# DNA modifications impact natural transformation of *Acinetobacter baumannii*

**DOI:** 10.1101/2023.02.09.527895

**Authors:** Nina Vesel, Christian Iseli, Nicolas Guex, Alexandre Lemopoulos, Melanie Blokesch

**Affiliations:** Laboratory of Molecular Microbiology, Global Health Institute, School of Life Sciences, Ecole Polytechnique Fédérale de Lausanne (EPFL), CH-1015 Lausanne, Switzerland; Bioinformatics Competence Center, Ecole Polytechnique Fédérale de Lausanne, CH-1015 Lausanne, Switzerland; Bioinformatics Competence Center, University of Lausanne, CH-1015 Lausanne, Switzerland

**Keywords:** *Acinetobacter baumannii*, natural competence for transformation, epigenome, restriction modification system

## Abstract

*Acinetobacter baumannii* is a dangerous nosocomial pathogen, especially due to its ability to rapidly acquire new genetic traits, including antibiotic resistance genes (ARG). In *A. baumannii*, natural competence for transformation, one of the primary modes of horizontal gene transfer (HGT), is thought to contribute to ARG acquisition and has therefore been intensively studied. However, knowledge regarding the potential role of epigenetic DNA modification(s) on this process remains lacking. Here, we demonstrate that the methylome pattern of diverse *A. baumannii* strains differs substantially and that these epigenetic marks influence the fate of transforming DNA. Specifically, we describe a methylome-dependent phenomenon that impacts intra- and inter-species DNA exchange by the competent *A. baumannii* strain A118. We go on to identify and characterize an A118-specific restriction-modification (RM) system that impairs transformation when the incoming DNA lacks a specific methylation signature. Collectively, our work contributes towards a more holistic understanding of HGT in this organism and may also aid future endeavors towards tackling the spread of novel ARGs. In particular, our results suggest that DNA exchanges between bacteria that share similar epigenomes are favored and could therefore guide future research into identifying the reservoir(s) of dangerous genetic traits for this multi-drug resistant pathogen.

## INTRODUCTION

The ability of bacteria to exchange genetic features by horizontal gene transfer (HGT), both within and between species, can drive rapid bacterial evolution. Alarmingly, this can result in the accumulation of dangerous genetic traits and the emergence of extremely problematic pathogens. Indeed, this is the case for the bacterium *Acinetobacter baumannii*, an ESKAPE pathogen, which is often able to escape treatment with multiple clinically-important antibiotics, and has therefore proven extremely problematic in hospital settings (1). *A. baumannii* was shown to exchange genetic information through HGT, but the relative contribution of conjugation, transduction, natural transformation, and other modes of HGT to resistance towards diverse antibiotics is not well understood (2). Nevertheless, since most of the antimicrobial (AMR) resistance genes in *A. baumannii* are located on the chromosome (3–5), natural transformation could potentially explain how most of these genes were acquired.

Natural competence for transformation is one of the primary modes of HGT (6). It is a physiological state in which the bacterium is able to take up DNA from the environment and either integrate it into its chromosome by homologous recombination or reconstitute it into a plasmid (7, 8). The DNA uptake machinery is conserved in naturally competent Gramnegative bacteria and involves the uptake of extracellular DNA into the periplasm via the retraction of a type IV pilus (T4P) (8–10). The incoming DNA is then bound by the DNA-binding protein ComEA, which acts as a Brownian ratchet, facilitating the entry of long pieces of DNA into the periplasmic space (10–12). In the next step, ComEC is thought to degrade one strand of the incoming DNA and, together with ComF, transfer the single stranded DNA (ssDNA) into the cytoplasm (8, 13). There, ssDNA is protected by Ssb and DprA, which also facilitates the loading of RecA (14). Finally, if there is sufficient homology, ssDNA is integrated into the host chromosome via the action of RecA and ComM, or, in case of plasmid DNA, the plasmid is reconstituted either by annealing of two complementary imported ssDNA strands or by the synthesis of the second DNA strand prior to circularization (8, 15–20).

To date, natural transformation has been experimentally demonstrated in several strains of *A. baumannii* (21–24). However, these studies reported considerable variation in the transformation frequencies obtained in the different strains. Some strains were non-transformable, while others were transformable with frequencies ranging from 10^-8^ to 10^-2^ (22–24). Similar observations have also been reported for the naturally competent bacterial species *Streptococcus pneumoniae* and *Haemophilus influenzae* (25, 26). While such variation in transformability might be a result of strain-specific competence regulation, it is also possible that DNA-specific factors could be involved. Furthermore, some studies have examined the transformability of *A. baumannii* using different kinds of donor DNA, including genomic DNA (gDNA), integrative or replicative plasmids, and PCR-amplified DNA fragments. While one study suggested that natural transformation occurs independently of the type of donor DNA used (27), another reported that transformation occurred most efficiently with purified gDNA, followed by integrative plasmids, and was the least efficient with PCR fragments (22). The increased efficiency of transformation with gDNA could be explained by longer regions of marker-flanking homology, yet the differences observed between the integrative plasmid and PCR product hint towards the involvement of DNA-specific features. Such features might be dependent on the source of the transforming DNA and should therefore be considered as a possible contributor to the observed variability.

Several DNA features have previously been shown to play a role in the transformability of certain naturally competent bacteria. Firstly, the sequence divergence between the transforming DNA and the acceptor’s chromosome was demonstrated to significantly diminish the bacterium’s natural transformability (28, 29). Secondly, it was reported that the presence of specific nucleotide sequences within the transforming DNA positively influences DNA uptake in the genera *Neisseria* and *Haemophilus*. These so-called DNA uptake sequences (DUSs) are conserved 9-12 bp long DNA motifs, which were shown to enhance the exchange of DNA between bacterial relatives belonging to the same species (30–33). Thirdly, the presence of DNA modifications was shown to affect transformability in diverse bacteria, such as *Helicobacter pylori, Campylobacter jejuni* and others (34–39). For instance, it was reported that a specific methylation motif, the result of a *Campylobacter-specific* orphan methylase, promotes DNA uptake in this organism (34). Importantly, DNA methylation patterns can also be a result of restriction modification (RM) systems. RM systems typically consist of a methylase and a cognate restriction enzyme that both recognize the same DNA sequence motif (40). Methylation protects the self-DNA from degradation by the restriction enzyme, while non-methylated foreign DNA is efficiently degraded (41, 42). The co-occurrence of RM systems and natural competence was demonstrated to affect transformability in *H. pylori*, *Pseudomonas stutzeri*, *C. jejuni* and *Neisseria meningitis* (35, 36, 38, 39). However, studies in *Bacillus subtilis* and *S. pneumoniae* reported only a minimal or no effect of RM systems on transformation (43, 44). Nonetheless, little is known about the impact of specific RM systems or DNA modifications on HGT in the genus *Acinetobacter*.

In this study, we aimed to understand how the source of the transforming DNA affects transformability in *A. baumannii*. This is especially relevant, since novel genetic traits such as antibiotic resistance genes (ARGs) are usually acquired from non-self-organisms. To unravel the involvement of specific DNA modifications in transformation, we used bacterial genetics and whole genome PacBio SMRT sequencing to identify specific genetic determinants responsible for the observed source DNA-dependent transformation phenotype. Additionally, we went on to investigate how the observed DNA selectivity is regulated, to understand whether its impact could be avoided under specific conditions. Collectively, our work highlights the importance of epigenetic modifications for natural transformability in the opportunistic pathogen *A. baumannii*.

## MATERIAL AND METHODS

### Bacterial strains, plasmids and growth conditions

The bacterial strains used in this study are listed in Table S1. The strains were cultured in lysogeny broth medium (LB; 1% tryptone, 0.5% yeast extract, 1% sodium chloride; Carl Roth, Germany) under shaking conditions (180 rpm) at 37 °C or on solid LB agar plates (1.5% agar) at 37°C. To select for *A. baumannii* after bi- or triparental mating with *E. coli* donor strains, CHROMagar *Acinetobacter* plates served as medium. These plates were prepared following the protocol provided by the manufacturer (CHROMagar, France). Gentamicin was added to the plates to select the colonies that acquired the respective transposon and apramycin was added to CHROMagar *Acinetobacter* plates to select for the bacteria carrying the envisioned (suicide) plasmids.

Transformation medium was prepared as described previously (45). Briefly, medium containing 5 g/L of tryptone, 2.5 g/L NaCl, and 2% agarose D3 (Euromedex, France; agarose from other manufacturers was non-functional for these assays) was autoclaved and kept at 60°C in liquid form. Prior to the transformation experiment(s), 1mL of this transformation medium was poured into 1.5 mL Eppendorf tube(s) and kept at room temperature (RT) to solidify.

To induce the P_BAD_ promoter, L-arabinose was added to the bacterial culture at the final concentration of 2%. Antibiotics were used at following final concentrations: kanamycin (50 *μ*g/mL), apramycin (100 *μ*g/mL), gentamicin (15 *μ*g/mL), ampicillin (100 *μ*g/mL).

### Genomic DNA isolation, PacBio Single Molecule, Real-Time (SMRT) sequencing and *de novo* genome assembly

Genomic DNA (gDNA) was purified as described previously (46, 47). Briefly, the *A. baumannii* strains ATCC17978, ATCC19606, and AB5075 were back-diluted 1:100 into 25 mL of LB medium and grown in an incubator under shaking conditions (180 rpm at 37°C) for 4 hours to reach an optical density at 600 nm (OD_600_) of 2-2.5 (for AB5075) or 1-1.5 (for ATCC17978 and ATCC19606). 15 mL of each culture was centrifuged to harvest the bacterial cells. To isolate gDNA from strains 29D2, 86II/2C, A118 (reSEQ), A118ΔH0N27_10820-30::kanR, A118ΔH0N27_10820::kanR-H0N27_10830-K176N, A118ΔH0N27_10830::kanR, A118ΔH0N27_10825::kanR, and ATCC17978-TnH0N27_12600, bacterial cells within 2 mL of an overnight culture were harvested by centrifugation. gDNA was isolated from all samples, using the Qiagen Genomic DNA buffer kit and 500/G (ATCC17978, ATCC19606, AB5075, AB5075-T, 29D2, 86II/2C) or 100/G (all other strains) columns, following the manufacturer’s instructions. The precipitated DNA was dissolved in 10mM Tris-HCl (pH 8.0). PacBio Single Molecule, Real-Time (SMRT) sequencing and *de novo* genome assemblies were performed at the Genomic Technology Facility of the University of Lausanne as previously described (47). The assembled genomes of all strains, except for A118 (reSEQ), were submitted to NCBI database and annotated with the Prokaryotic Genome Annotation pipeline (PGAP). The sequencing statistics of all the strains submitted to NCBI and the NCBI GenBank accession numbers are shown in Tables S2 and S3.

### Analysis of DNA modification motifs

DNA modifications were determined based on the SMRT sequencing results of this study plus the output obtained for strain A118 for which we previously reported its genome assembly (46). The raw data of strains A118, ATCC17978, and ATCC19606 were analyzed using the smrtlink-release_7.0.1.66975 software suite, while the raw data from all other sequenced strains were analyzed using the smrtlink-release_10.2.0.133434 software suite. Both software suites were obtained directly from the vendor’s webpage: https://www.pacb.com/support/software-downloads/. The tool named “Base Modification Analysis Application” was used to detect modified DNA bases in the bacterial genomes, and the corresponding DNA motifs surrounding modified bases, as outlined in this White Paper: https://www.pacb.com/wp-content/uploads/2015/09/WP_Detecting_DNA_Base_Modifications_Using_SMRT_Sequencing.pdf. The sequencing reads of the different data sets were mapped onto the corresponding reference genome sequences of each strain and analyzed using the default parameters of the tool (95% confidence interval bounds, P-value cutoff = 0.001, minimum methylation fraction =0.3 and minimum Qmod score = 100) and activating its “Find Modified Base Motifs” option. For each analyzed sample, the tool produced a table listing the genomic positions at which a base modification was detected, as evidenced by the deviation of the inter-pulse duration from the expected base incorporation time. The tool’s output listed motifs that are present in more than 30% of the occurrence of that motif around detected modified bases. All motifs identified in at least one sample were collected and a perl script was written to count the number of occurrences of each motif in the whole genome of each sample, and to compute the fraction of each motif that contained the detected modified bases. The modified positions were represented on Circos plots using the R package circlize (48), and a summary heatmap comparing the fraction of modified motifs found in each sample was produced using the R function heatmap.2 from the library gplots package 3.1.3 (49).

### Identification of potential methylases in different *A. baumannii* isolates

To investigate the potential methylases across *A. baumannii* isolates, we performed a pangenome reconstruction using the PPanGGOLiN (v. 1.2.74) pipeline (50) (all options with default parameters; both nucleotide sequences and annotation files were used as inputs to the pipeline). Briefly, PPanGGOLiN uses genome sequences and annotations to reconstruct pangenomes and subsequently build gene-family based partitions. The program integrates information from the annotation (protein coding genes, CDS, genomic neighborhood) to define gene families (for more details on the method, see (50)). Next, genes are split into persistent and cloud families. PPanGGOLiN identified 5181 genes (3108 persistent and 2073 cloud genes) across the 6 strains. Out of the obtained gene repertoire, we identified 111 genes which contained “methyl” in their annotation, and, after individual curation, we singled out 29 specific DNA methylase / methyltransferase genes. Based on their presence / absence in each strain, we reconstructed a heatmap in R environment (v 4.2.1) (51) using the pheatmap package v 1.0.12 (52) (default parameters with no clustering).

### Comparison of RMC-containing A118 genomic region with other *A. baumannii* strains

Genome alignment of strains A118, ACC17978, ATCC19606, AB5075, 29D2, and 86II/2C was performed in Geneious Prime (v 2022.2.2) with the progressiveMauve algorithm (53). The genomic region carrying the methylase H0N27_10820 within the RMC in strain A118 was compared with the same genomic region in other strains to identify conserved upstream and downstream flanking genes. After obtaining the specific sequences and annotations for this genomic region in the six different strains, we aligned, compared, and visualized the region using Clinker (v 0.0.25) (54) with default options.

### Construction of mutants in *A. baumannii*

Most of *A. baumannii* mutants (see Table S1) were constructed following a natural transformation protocol as described previously (45) with few modifications. To genetically engineer strains, PCR amplification products served as transforming material. These PCR fragments were amplified using PWO polymerase (Roche) or Q5 polymerase (New England BioLabs). Each fragment consisted of an antibiotic resistance cassette (kanamycin or gentamicin resistance genes encoded by *aph* and *aacC1*, respectively) flanked by 1-kb long regions homologous to the desired chromosomal insertion locus. The final PCR fragments were generated by overlapping PCR using the three individual fragments as template, and were added to respective bacterial strains, as described below for the natural transformation assay. The transformed cells were resuspended in PBS and plated on selective LB agar plates (containing kanamycin or gentamicin). Selected transformants were screened by colony PCR using GoTaq polymerase (Promega) and the correct DNA sequences were confirmed by Sanger sequencing (Microsynth, Switzerland).

To obtain a mutant in non-transformable strain ATCC19606, a standard suicide plasmid-based mating approach was used, as described previously (46). Briefly, a derivative of the counter-selectable plasmid pGP704-Sac28 (55), carrying the desired deletion construct was engineered by standard molecular methods. The two flanking regions of the chosen genomic exchange region were amplified and joined with PCR fragment carrying antibiotic resistance cassette by overlapping PCR using Q5 polymerase (New England BioLabs). Next, this final PCR fragment was cloned into plasmid pGP704-Sac28. The cloned construct was transferred into *E. coli* strain S17-1λpir and confirmed by colony PCR (GoTaq polymerase, Promega) and Sanger sequencing (Microsynth, Switzerland). The correct plasmid was transferred to *A. baumannii* by biparental mating. Subsequent sucrose-based counter-selection was used to select plasmid-cured clones after a putative allelic exchange event as previously described (46, 55). Allelic exchanges were confirmed by Sanger sequencing (Microsynth, Switzerland).

To electroporate the plasmid pABA, a 2 mL of overnight culture of the respective strain was centrifuged for 10 min at 10’000xg at room temperature (RT). The pelleted bacteria were washed twice with 1 mL of sterile 10% glycerol and resuspended in 100 μL of 10% glycerol. 2 μL of plasmid (400 ng) was added to 50 μL of such electrocompetent cells and electroporated inside an electroporation cuvette at 1.8 kV. Immediately after the electroshock, 950 μL of 2xYT was added to the bacteria to foster their recovery. The suspension was carefully transferred into 14 mL round-bottom tubes and grown aerobically for 1 hour at 37°C. After incubation, the cells were spread onto LB agar plates containing apramycin and incubated at 37°C overnight. The presence of the plasmid was confirmed by colony PCR (GoTaq polymerase, Promega). For construction of ATCC17978Δhcp::aprR, a PCR product with an amplified AprR cassette flanked by 1-kb long regions homologous to the desired chromosomal insertion locus was used instead of the plasmid during electroporation.

To construct mutants with inducible genes, the respective genes were cloned behind the arabinose-inducible promoter P_BAD_ located on the miniTn7 transposon (=TnAraC), which itself was carried on the suicide plasmid pGP704 (e.g., plasmid pGP704-TnAraC; (47, 56)). The engineered plasmids were transferred into *A. baumannii* through standard triparental mating (57), which, assisted by a T7 transposition machinery-encoding helper plasmid, resulted in the transposition of the Tn-AraC-XX (abbreviated as TnXX) construct downstream of the *glmS* gene on the *A. baumannii* chromosome.

### Natural transformation assay

Natural transformation assays were performed as previously described (45) with minor modifications. Briefly, overnight cultures of *A. baumannii* were back-diluted 1:100 in 2 mL LB medium and grown aerobically at 37°C until they reached an OD600 of approximately 1. Next, the cultures were diluted 1:100 in sterile PBS and 2 μL of this bacterial suspension was mixed with the same volume of transforming DNA at a concentration of 50 ng/μL (for pABA) or 100 ng/μL (for gDNA or PCR products). 3 μL of the mixture was subsequently spotted onto 1 mL of solidified transformation medium inside a 1.5 mL Eppendorf tube and incubated at 37°C for 24 hours. The next day, 300 μL of PBS was added and the Eppendorf tube vortexed at max. speed to resuspend the cells. The bacterial suspension was serially diluted and spotted in duplicate on agar plates containing the appropriate antibiotic (apramycin or rifampicin) to select for transformants. Dilutions were also spotted onto plain LB agar plates to estimate the total number of bacteria. After overnight incubation at 37°C, transformation frequencies were calculated as the number of colony forming units (CFU) of the transformants divided by total number of CFUs. Data in graphs represent the average of three biologically independent replicates. When no transformants were detected, the values were set to the detection limit to allow calculation of the average and for statistical analysis using the software Prism (GraphPad; San Diego USA). For statistical analysis, transformation frequencies were log-transformed (58). Detailed specifications of statistical analyses are provided in the figure legends.

The gDNA samples that were used as transforming material in the transformation assays were purified using the Qiagen Genomic DNA buffer kit and Qiagen 100/G columns. Purification was based on 2 mL of overnight-grown cultures, as described above, and the precipitated gDNA was dissolved in 10 mM Tris-HCl (pH 8.0) buffer.

The pABA plasmid was purified using PureYield^™^ Plasmid Miniprep System (Promega), starting with 5 mL of an overnight grown culture. When indicated for gene induction, overnight cultures were grown with 2% arabinose. When specified, the pABA plasmid (from *E. coli* INV110) was methylated with Dam methylase (New England BioLabs), following the manufacturer’s instructions.

PCR products that served as transforming material were obtained by amplification with PWO polymerase (Roche) or Q5 polymerase (New England BioLabs).

### Quantitative reverse transcription-PCR (qRT-PCR)

Overnight cultures were back-diluted 1:100 into 2 mL LB medium (in duplicate for the 2-hours samples) and grown aerobically for two or six hours (as specified) at 37°C. Next, the cultures were centrifuged for 3 min at max speed and 4°C. Bacterial pellets were washed with 1 mL of sterile and cold PBS and resuspended in 1 mL of TRI Reagent (Sigma-Aldrich). After vortexing for 10 seconds, the samples were snap-frozen in a dry ice-ethanol bath and stored at −80°C. Subsequent steps included RNA extraction, DNase treatment, reverse transcription and quantitative PCR (qPCR), which were performed as described previously (59). Expression levels of individual genes were normalized to the transcript levels of the housekeeping gene *gyrA* (=relative expression) and the average was calculated from three biologically independent experiments. Relative expression values were log-transformed (58) for statistical analysis using the Prism software (GraphPad; San Diego USA). The parameters of the statistical analyses are described in the figure legends.

## RESULTS AND DISCUSSION

### Source of transforming DNA affects transformation in *A. baumannii*

To better understand how efficiently different *A. baumannii* isolates can acquire DNA by natural transformation, we compared the transformation frequency of four *A. baumannii* strains, using gDNA originating from six different *A. baumannii* strains (Table S4). Specifically, strains A118, AB5075, 29D2 and 86II/2C served as DNA acceptors, since they were previously shown to be naturally competent (21, 22, 45). The donor gDNA, used as the transforming material, was purified from an apramycin cassette-containing (AprR) derivative of each of the four transformable strains mentioned above, as well as two non-transformable strains, ATCC17978 and ATCC19606 (Table S4). To obtain AprR-marked donor strains, the apramycin resistance cassette was inserted into the *hcp* gene of each strain. We chose *hcp*, which encodes a part of the type VI secretion system (T6SS), as a neutral locus, given that the T6SS is not involved in competence development or DNA uptake. Importantly, the T6SS gene cluster is highly conserved in the different *A. baumannii* strains, which should facilitate homologous recombination during transformation. Indeed, the pairwise identity of the genomic region 10 kb upstream and downstream of *hcp* is higher than 95% for most of the compared strains (Fig. S1A). In contrast, strain ATCC17978 carries several insertions upstream of *hcp*, resulting in a significantly decreased pairwise identity of less than 47% when compared to the other tested strains (Fig. S1A). Nevertheless, the region of approximately 1 kb immediately upstream and downstream of *hcp* remains highly conserved in all strains, with only a few single nucleotide polymorphisms (SNPs) when compared to strain A118, which served as the reference strain in this study (Fig. S1B).

Since the *hcp* chromosomal region is highly homologous between the tested strains – at least in close proximity to the AprR cassette – one might expect that the transformation frequencies should be similar when using gDNA from different sources. Interestingly, however, we observed several significant differences in transformation frequencies in all tested acceptor strains when the donor gDNA had distinct origins (Fig. 1A). These differences appear to be mostly independent of sequence homology between the acceptor strain and the donor DNA. Strikingly, in strain A118 (Fig. 1B) transformation frequencies were 10- to 40-fold lower when the DNA originated from any of the non-self strains. Notably, even though the transformation frequency was the lowest (~40-fold reduction) with the ATCC17978 donor gDNA, this was only a small reduction compared to the other nonself gDNA sources, suggesting that the reduced pairwise identity between the two strains has only a minor negative effect. A slight decrease in transformation levels when using ATCC17978-derived gDNA in comparison to self gDNA was also observed when strains AB5075 and 29D2 served as DNA acceptors, whilst transformation frequencies were comparable for strain 86II/2C (Fig. 1A). A minor or no effect of sequence divergence on transformation frequency was somewhat expected, since recombination can occur between highly similar DNA segments of only 25 to 200 bp in length (21). Therefore, the highly similar regions either proximal or distant to the *hcp* gene might be sufficient to reach high level of transformation when gDNA from ATCC17978 was used as transforming material.

**Figure 1.**
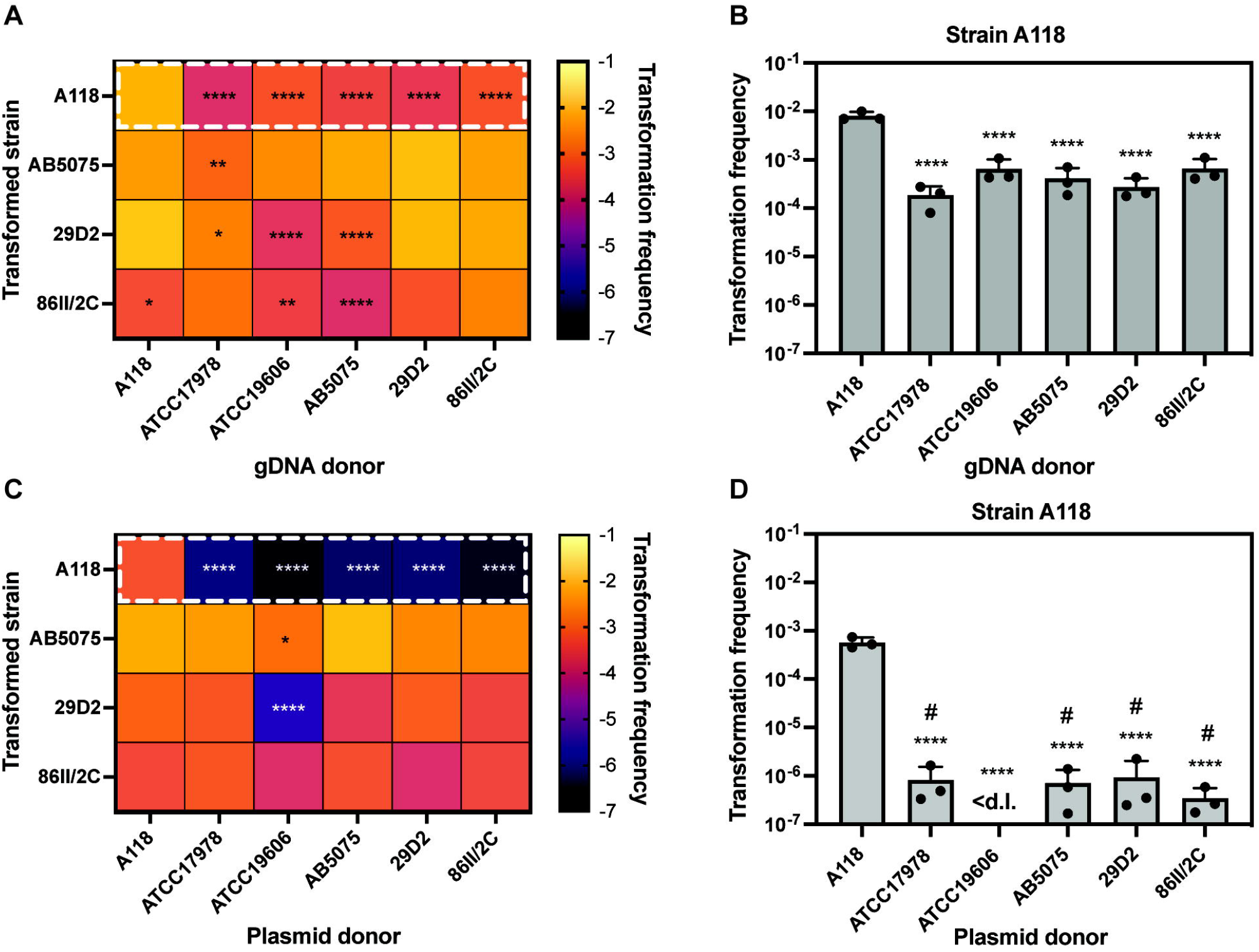
Origin of the transforming DNA impacts natural transformability. **(A)** Transformability of all transformable strains tested (indicated on *Y*-axis) using gDNA from different *A. baumannii* strains as transforming material (indicated on *X*-axis). The heat map indicates the transformation frequencies according to the legend. **(B)** Transformation frequency of strain A118 after transformation with genomic DNA (gDNA) originating from different *A. baumannii* strains. Details of boxed area in panel A. **(C)** Transformability of strains using plasmid pABA derived from the different *A. baumannii* strains as transforming material. Heatmap as in panel A. **(D)** Transformation frequency of strain A118 when transformed with plasmid DNA (pABA) originating from different *A. baumannii* strains. Details of boxed area in panel C. The mean values (± standard deviation, SD) of three independent biological replicates are shown for panels B and D with each individual experiment being represented as a dot. <d.l., below detection limit. #, below detection limit in at least one experiments. Statistics were performed on log-transformed data, using twoway analysis of variance (ANOVA) with Dunnett’s (A, C) or Tukey’s (B, D) correction for multiple comparisons. Only statistically significant differences are shown. *, *P* < 0.05, **, *P* < 0.01, ****, *P* < 0.0001.

To exclude any potential effects of sequence homology when investigating the impact of the source of DNA on transformation efficiency, we repeated the experiment using the small (6.8kb) replicative plasmid pABA as the donor DNA. pABA is a derivative of the conventional plasmid pBAD18-Kan, wherein an *Acinetobacter*-specific origin of replication has been added to enable replication in *A. baumannii* (60) and the KanR cassette (*aph*) has been replaced with an AprR cassette (*aac(3)IV*). This plasmid was electroporated into strains A118, ATCC17978, ATCC19606, AB5075, 29D2, and 86II/2C and subsequently repurified using standard plasmid mini-preparations. Next, we used these strain-passaged plasmids in a transformation assay with all the transformable strains as acceptors. Interestingly, we again observed significant DNA origin-dependent transformation differences in some acceptor strains (Fig. 1C). In strain A118 the DNA-source dependent selectivity was conserved irrespectively of the DNA type. Indeed, transformation frequencies with non-self-originating plasmids were barely above detection limit, while transformation with the plasmid originating from the strain A118 itself was around three orders of magnitude higher (Fig. 1D). For the other naturally transformable strains, the difference between the self- or non-self-originating plasmid was less pronounced, with the notable exception of strain 29D2 transformed with the plasmid isolated from ATCC19606 (Fig. 1C). Given the strong DNA source-dependent effect on transformation when using gDNA and plasmids, we hypothesized that epigenetic DNA modifications, might either enhance and/or limit natural transformability in some *A. baumannii* strains, such as A118. This could also be the reason for the discrepancies between DNA source-dependent transformation patterns when gDNA or plasmids were used as transformation material and could be explained by the absence of some specific DNA modification recognition sites on the pABA sequence (Table S5). This hypothesis is in line with previous work on *H. pylori, Pseudomonas stutzeri*, *C. jejuni* and *Neisseria meningitidis* where it was shown that methylated DNA motifs could affect transformation by either impacting the specificity of DNA uptake or the degradation of incoming DNA (34–36, 38, 39).

### SMRT sequencing reveals diverse DNA modification patterns among different *A. baumannii* strains

To understand whether the DNA source-dependent differences in transformation efficiency could be explained by strain-specific epigenomes, we whole genome sequenced all tested *A. baumannii* strains, except previously sequenced strain A118 (46), using PacBio SMRT sequencing (Table S2). This technique not only determines the genome sequence, but also provides information regarding various DNA modifications such as N^6^-methyl-adenine (m6A), N^4^-methyl-cytosine (m4C) and N^5^-methyl-cytosine (m5C) (even though the latter modification requires additional conversion steps pre-sequencing or very deep sequencing efforts), with single-nucleotide resolution (61). Interestingly, as shown in Figure 2A, all the sequenced strains exhibited a specific DNA modification pattern, mostly comprising several methylation motifs. For all strains, the identified DNA modifications belonged to 6-15 nucleotide long motifs, mostly carrying methylation marks on either adenine residues (8x) or on cytosine residues (1x). For 6 out of the 15 identified DNA modification motifs, only the modified residue could be determined, but not the type of the modification (labeled ‘unk’ for unknown in Figure 2A). Several of the identified DNA modification motifs were present in more than one strain, for example m6A:GAAAGC:4 (meaning a m6A modification on the 4^th^ nucleotide of this motif) was common to strains AB5075 and 29D2. Other modifications were strain-specific, such as m6A:RGATCY:3 in strain A118. We observed different proportions of modified recognition sites relative to their abundance within each genome, which varied from approximately 20 to 100 percent (shown on a scale from 0 to 1 in the legend of Figure 2A).

**Figure 2.**
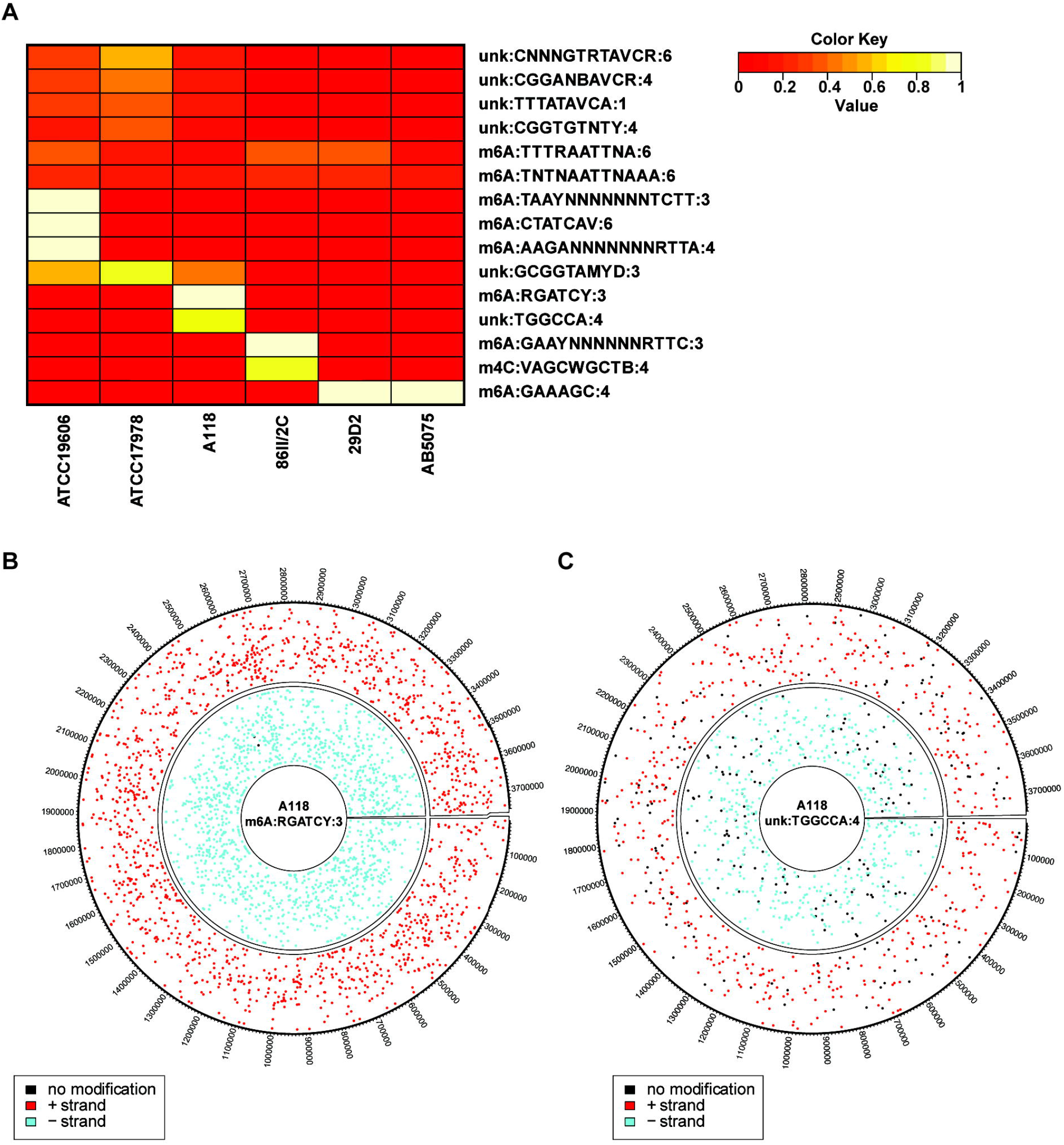
SMRT sequencing reveals DNA modifications in diverse *A. baumannii* strains. **(A)** The heatmap shows the abundance of the 15 DNA modification motifs, which were detected on one or several of the six different *A. baumannii* strains. The color indicates the proportion of modified recognition sites, as shown by the color key. unk, unknown type of modification. **(B-C)** Spatial distribution of the DNA modification m6A:RGATCY:3 **(B)** and unk:TGGCCA:4 **(C)** on both strands of the A118 chromosome. Modified recognition sites are shown in color (red and blue for + and – strand, respectively), while unmodified sites are shown in black.

To date little is known about the epigenome of the genus *Acinetobacter*. A study identified a methylase in *A. baumannii* strain ATCC17978 – AamA, that methylates adenine at GAATTC motifs and was suggested to be involved in surface-associated motility (62). Even though *aamA* homologs were present in all the tested strains (Fig. S2), we did not detect m6A:GAATTC:3 modification in any of them. However, two similar motifs - m6A:TTTRAATTNA:6 and m6A:TNTNAATTNAAA:6, were partially methylated in all the tested strains (Fig. 2) and could therefore be a result of the AamA methylase. Alternatively, AamA might not be produced under the tested conditions.

The sole report of a complete *Acinetobacter* methylome was performed in *Acinetobacter calcoaceticus* 65, for which several DNA modification motifs were identified, belonging to either type I or type II RM systems (63). There were no common DNA modification motifs between the reported methylome of *A. calcoaceticus* 65 and the methylomes of the *A. baumannii* strains determined in this study. This is not surprising given the high variability that we already observed in the modification profiles between the different isolates. Furthermore, as RM systems are dynamic and their genes are often acquired through HGT (28), such variability between members of the same genus or species is to be expected.

We considered two distinct hypotheses to explain how the variability of DNA modifications could impact the transformability of strain A118, whereby transformation frequency with self-DNA was higher than with non-self DNA. First, the presence of DNA modification on the donor DNA could lead to reduced transformation in A118 if, for instance, this DNA modification was recognized by a A118-specific type IV restriction enzyme. Type IV restriction enzymes, unlike the other types, degrade modified DNA (42). However, analysis of our modification data showed that no specific DNA modification was shared between all the tested strains, but absent in strain A118 (Fig. 2A). We therefore considered a second possible scenario whereby an A118-specific DNA modification could either promote DNA uptake or prevent DNA degradation. The latter could be explained by the presence of an A118-specific RM system that would result in methylation and therefore protection of self-DNA, while promoting the degradation of the non-self DNA lacking these specific methylation marks by the corresponding restriction enzyme. Interestingly, we identified two A118-specific DNA modification motifs – m6A:RGATCY:3 and unk:TGGCCA:4 [which we characterized as m4C-TGGCCA:4 in further sequencing runs; see below] – which, as depicted in Figure 2B-C, are randomly distributed throughout the entire genome. Since the recognition site TGGCCA is absent from the plasmid pABA (Table S5), we hypothesized that the m6A:RGATCY:3 DNA modification was likely responsible for the observed phenotype. As shown in Figure 2B, nearly 100% of the RGATCY recognition sites were methylated on both strands of the A118 chromosome, suggesting that the responsible methylase is highly active and could therefore be involved in efficient DNA protection.

### Comparing methylase repertoire between different *A. baumannii* isolates

To determine the possible candidates for m6A:RGATCY:3 DNA modification, we compared the annotated genomes of the six PacBio-sequenced *A. baumannii* strains. As shown in Figure S2, we identified a repertoire of genes encoding for proteins with DNA methylase-like activity in each tested strain. This enabled us to assess two A118-specific DNA methylases: H0N27_12600 and H0N27_10820 (protein names are based on their locus tags according to NCBI accession number CP059039), which could be responsible for the m6A:RGATCY:3 DNA modification. The H0N27_12600, is annotated as a DNA adenine methylase, but lacks an obvious cognate restriction enzyme(s) within its gene neighborhood. Thus, while we cannot exclude that a partner restriction enzyme might be encoded elsewhere on the genome, we consider H0N27_12600 to be an orphan methylase [referred to as MT2 throughout the text], which is unlikely to be responsible for the observed transformation phenotype. The second methylase H0N27_10820 is also annotated as DNA adenine methylase and is located close to a restriction endonuclease-encoding gene (H0N27_10830; Fig. 3A). This suggested that the methylase H0N27_10820 might be a part of type II restriction modification (RM) system, which are often involved in defense against foreign DNA (42). We therefore aimed to further characterize this putative RM system and the genes located in its neighborhood.

**Figure 3.**
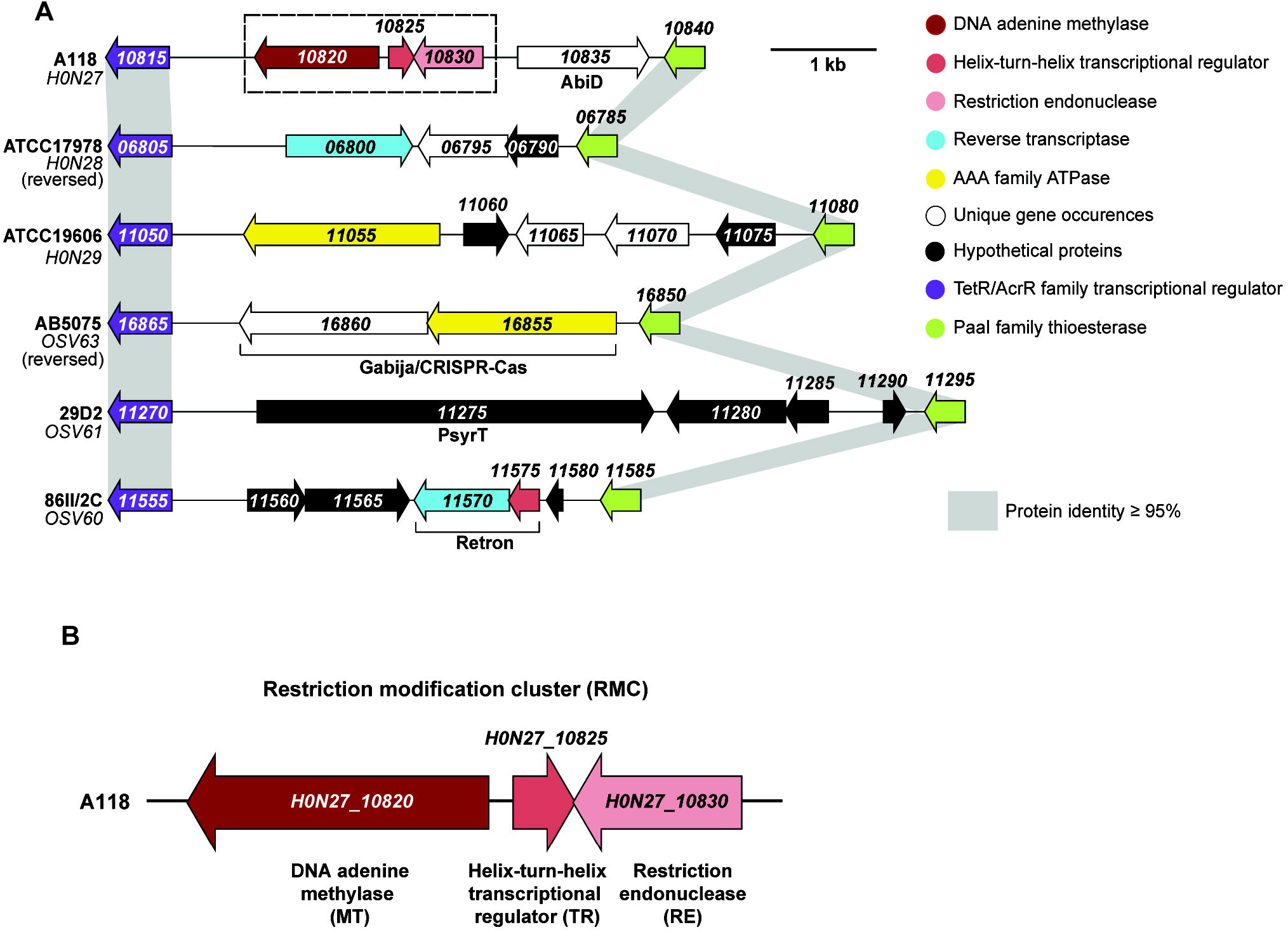
Strain A118 carries unique RM system. **(A)** Comparison of A118 genomic region carrying the gene encoding the predicted methylase H0N27_10820 with the same region of other *A. baumannii* strains. Predicted annotations for gene products are represented with different colors, as shown in the legend. Genes that encode for defense systems according to predictions by PADLOC or DefenseFinder are defined underneath the gene number(s). **(B)** Zoom of the A118-specific genomic region, carrying the newly identified RM system genes *H0N27_10820-30 (H0N27_10820* (MT) in dark red, *H0N27_10825* (TR) in red and *H0N27_10830* (RE) in light red).

Interestingly, the potential RM system is encoded in a region of the genome that exhibits high genetic variability between different strains (Fig. 3A), suggesting that it might be a hot-spot for genomic integration. Given the involvement of RM systems in bacterial defense against foreign DNA (42) we wondered whether other *A. baumannii* strains also carried genes encoding DNA defense systems in the same region, potentially forming a small defense island. We therefore inspected this region in our tested strains using the Prokaryotic Antiviral Defence LOCator (PADLOC) (64) and DefenseFinder tools (65, 66). This approach identified several genes encoding for known anti-phage defense systems such as: Gabija/CRISPR-cas complex in strain AB5075, *psyrT* - belonging to a toxin-antitoxin system in strain 29D2, and a retron defense unit in strain 86II/2C (Fig. 3A). Strains ATCC17978 and ATCC19606 also harbored genes encoding for a reverse-transcriptase and an AAA family ATPase, respectively, which are often associated with bacterial DNA defense (67–69). The genomic region specific to strain A118 encompasses the four genes H0N27_10820 to H0N27_10835, coding for the aforementioned RM system (DNA adenine methylase H0N27_10820 [MT] & restriction endonuclease H0N27_10830 [RE]) interspersed by a helix-turn-helix transcriptional regulator [TR]-encoding gene H0N27_10825 and followed by gene H0N27_10835, which was predicted by DefenseFinder to encode for AbiD, involved in abortive infection during anti-bacteriophage defense (Fig. 3A). Protein BLAST analysis of H0N27_10830 suggests that the putative RE belongs to the Endonuclease *BglII/BstYI* superfamily, which typically recognizes AGATCT or RGATCY. This strongly suggested that this RM system is likely responsible for A118-specific DNA modification m6A:RGATCY:3 and ultimately could explain the DNA source-dependent transformation phenotype.

### The A118-specific RM system impacts the strain’s natural transformability

To determine whether the H0N27_10820-30 RM system (Fig. 3B) (henceforth referred to as RMC = restriction modification cluster) is indeed responsible for the DNA origin-dependent differences in transformability of strain A118, we designed a RE-deficient strain A118ΔH0N27_10830 (ΔRE) and compared transformation frequencies in the WT strain and the RE-deficient strain, using various types of donor DNA. Firstly, we used gDNA derived from different strains. As expected, no significant difference was observed when using self-DNA from A118 (Fig. 4A). Importantly, however, when comparing the transformation frequencies of the RE-minus mutant to those of the WT strain, we observed a significant 20 to 40-fold increase in transformation with the gDNA samples that were derived from the foreign strains representing non-self DNA. Secondly, when we used a PCR amplified fragment of Δ*hcp*::AprR as the donor DNA, which contains 1 kb flanking regions on either side and lacks any DNA modifications, we reproducibly observed a slight, but significant increase in transformation in the RE-deficient mutant (Fig. 4B). Thirdly, when plasmid pABA originating from different strains was used as the donor DNA, the DNA-origin effect was almost completely abolished in the RE-deficient strain (Fig. 4C). The DNA motif RGATCY, carrying methylation marks, likely responsible for DNA selectivity, was present in all the donor DNA types (Table S5). These results strongly suggest that the RE limits acquisition of foreign DNA independently of transforming DNA type. Interestingly, the effect of the RE on transformation was the least drastic for the PCR product, more pronounced for the diverse genomic DNA samples, and strongest when plasmids were used as transforming material. These differences might be partially due to the different copy number of the resistance gene when using distinct DNA types, which was the highest for the PCR product and the lowest for the gDNA. While this could explain the modest RE-dependent effect observed with the PCR product, the severity of the RE impact on transformation with plasmid DNA might be rather due to the different mechanisms that are required to establish plasmid or linear homologous DNA in the recipient cell.

**Figure 4.**
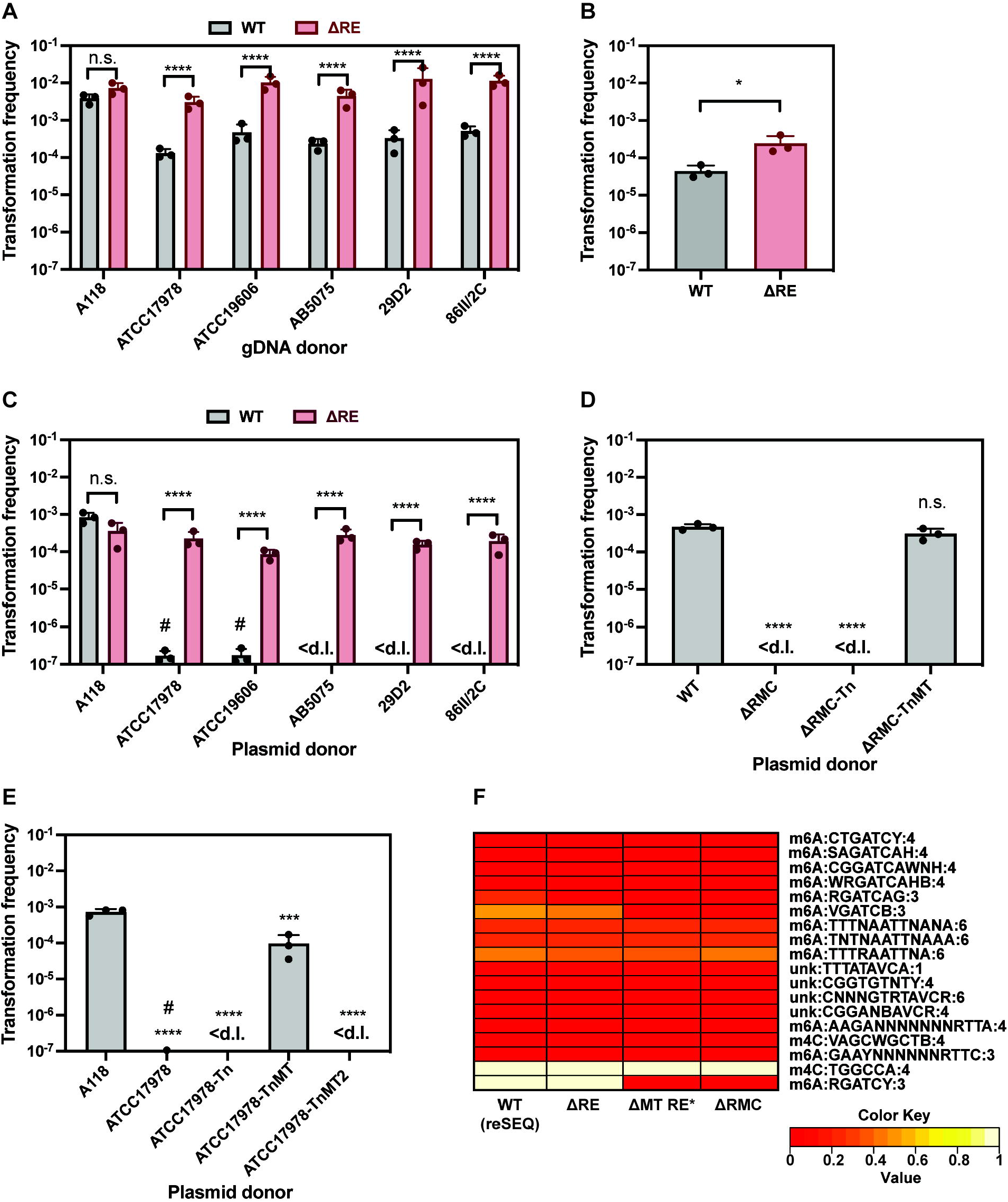
The RMC is responsible for source DNA-dependent transformability of A118. Transformability of strain A118 (WT) and A118ΔH0N27_10830 (ΔRE) using **(A)** gDNA of different *A. baumannii* strains, **(B)** PCR product amplifying *Δhcp::aprR* region, or **(C)**plasmid pABA originating from different strains as transforming material. **(D-E)** Transformation of strain A118 with plasmid pABA originating from various strains grown with 2% L-arabinose: **(D)** A118 (WT), ΔRMC (A118ΔH0N27_10820-30), ΔRMC-Tn (ΔRMC carrying transposon control), or ΔRMC-TnMT (ΔRMC complemented with inducible H0N27_10820 (MT) on transposon); **(E)** A118, ATCC17978, ATCC17978 carrying empty transposon (Tn) or transposon with inducible copy of H0N27_10820 (TnMT) or H0N27_12600 (TnMT2). **(F)** The RMC cluster leads to m6A modified motif RGATCY. Heatmap showing the abundance of different DNA modification motifs in the RMC-related mutants of strain A118. Details as described for Figure 2. The average values (± SD) of three independent biological replicates are shown (A-E). <d.l., below detection limit. #, below detection limit in at least one replicate. Statistical analyses were performed on log-transformed data using a two-way ANOVA with Šidák’s correction for multiple comparisons (A, C), an unpaired t test (B), or a one-way ANOVA with Dunnett’s correction for multiple comparisons (D, E). *, *P* < 0.05, ***, *P* < 0.001, ****, *P* < 0.0001; n.s., not significant.

All types of transforming DNA enter the cell in a single stranded form, which is believed to be inaccessible to most restriction endonucleases, which instead act on dsDNA (70). Furthermore, the incoming ssDNA is thought to be protected from degradation through the tight binding of the Ssb and DprA proteins (14, 71). Upon entry into the cell, gDNA and PCR products are integrated into the genome via homologous recombination. The reconstitution of replicative plasmids, on the other hand, requires either import and annealing of two complementary ssDNA strands or the synthesis of the second strand, prior to circularization (19, 20). Therefore, plasmid cleavage most probably occurs immediately after dsDNA is formed / synthesized and before it is protected by methylation. In contrast, it is not well understood at which integration step the linear transforming DNA is cleaved by restriction endonuclease, even though RM-dependent decrease in transformation with integrative DNA was also demonstrated in some other bacterial species (35, 38, 39). It was recently shown that the heteroduplex is resolved by replication during homologous recombination (72, 73). Given that methylases act on hemi-methylated and non-methylated DNA, while restriction endonuclease can only cleave non-methylated DNA, it was suggested by Johnston *et al*. that upon replication, homologous DNA stretches of integrative DNA result in hemi-methylated dsDNA, while heterologous DNA stretches become unmethylated dsDNA regions. Such regions represent an ideal substrate for degradation, resulting in death of the transformed cells (74). To test whether this could explain the effect of the RE on integrative DNA in *A. baumannii*, we used a fully homologous PCR product carrying a SNP in *rpoB* (conferring rifampicin resistance) as the donor DNA. As shown in Figure S3A, the RE had no effect on transformation frequency under these conditions, despite RGATCY recognition site was present on the PCR product serving as transforming DNA (Table S5). This suggests that the presence of heterologous region on the integrative DNA is required for the RE-dependent effect on transformation, which was also observed in *S. pneumoniae* and *N. meningitidis* (39, 75). While our data indicate that RM systems reduce acquisition of novel genes, it is reasonable to speculate that the transformation-driven loss of certain genetic regions might not be impacted.

Since the RE exclusively impaired transformation with non-self-derived DNA, the logical assumption is that the MT protects self-DNA from the RE-mediated degradation by adding specific methylation marks. To test this assumption, we attempted to delete both the entire RMC gene cluster and the MT gene alone. However, in further support of a functional connection between the MT and RE, it was impossible to delete the MT alone, except in the presence of a spontaneous suppressor mutation in the RE (further referred to as ΔMT RE* strain). This is likely because in the absence of the methylation motifs, the host genome would become susceptible to degradation by the RE. Next, we purified plasmids from the RMC- or MT-minus strains and used them as donor DNA for the WT strain of A118. As shown in Figures 4D and S3B, the transformation frequency decreased significantly when the plasmid originated from the strains lacking either the RMC (ΔRMC) or the MT (ΔMT RE*), confirming the protective effect of this specific methylation on the transforming material if derived from WT cells. In contrast, plasmids originating from the methylase-deficient strains that were complemented by an arabinose-inducible MT gene *in cis* (Tn-MT) were able to support transformation at levels comparable to the WT control (Figs. 4D and S3B).

Additionally, to determine whether the expression of the A118-specific MT in a different strain background could increase the transformation potential of non-self-derived DNA, we introduced the arabinose-inducible TnMT construct into the strain ATCC17978. By doing so, the transformation frequency was significantly increased when we used plasmids derived from this strain (grown under MT-inducing conditions) as compared to either the parental strain or an empty transposon-carrying control (Fig. 4E). Interestingly, while increased by several orders of magnitude compared to the controls, the transformation efficiency did not reach the same level as with A118-derived plasmid DNA, which could be due to the presence of additional DNA modifications in this strain.

In contrast to the results above, no transformants were obtained with plasmid DNA purified from ATCC17978 expressing the arabinose-inducible A118-specifc orphan methylase H0N27_12600 (TnMT2; Fig. 4E). This result suggests that MT-2 does not play a DNA-protective role during natural transformation. However, the absence of protection could also be linked to the fact that the gene’s induction did not work properly in our experiments or that the methylase was non-functional. To address these possibilities, we SMRT sequenced strain ATCC17978-TnMT2 after arabinose-dependent induction of the MT2 gene. The results showed that the induction of MT2 resulted in the additional DNA modification - m4C:TGGCCA:4 (Fig. S3C). Importantly, this DNA modification was also present in strain A118 (Fig. 2A), yet the exact modification (methylation) only became apparent in additional sequencing runs. This therefore showed that MT2 is responsible for the m4C:TGGCCA:4 DNA modification but that it does not have a protective role against the RE. Collectively, our results suggest that the RMC cluster-encoded MT is the only methylase that is responsible for efficient DNA protection in the RE-producing strain A118.

Finally, to confirm that the m6A:RGATCY:3 methylation is a result of the identified RMC, we performed PacBio SMRT sequencing to assess the DNA methylome of the RMC-related mutants of strain A118 (Table S3). As a control, we re-sequenced the WT A118 strain, to which we refer as WT(reSEQ) (previously sequenced version of strain A118 referred as WT). The DNA methylomes of both sequencing versions of the WT were slightly different, which might be explained by the differences in gDNA preparation and sequencing machines (see Methods). However, as shown in Figure 4F, the deletion of the RE did not affect DNA modifications, as they were almost identical to those of the WT(reSEQ). In contrast, the DNA modification m6A:RGATCY:3 was no longer identified in the strains lacking either the entire RMC or the MT plus the RE* (Fig. 4F). This confirmed that the identified DNA modification was indeed the result of the MT within the RMC. Interestingly, Nucleotide BLAST analysis of the A118 RMC identified similar or identical RM systems in several *A. baumannii* strains, including Ab42, UH6507, ARLG_1772 *etc*. as well as in other members of the *Acinetobacter* genus (Table S6).

### A helix-turn-helix regulator controls methylase expression

Regulation of RM systems frequently relies on conserved helix-turn-helix transcriptional regulators, which control the homeostasis between the restriction endonuclease and the methylase (76–80). Since the RMC also carried the same type of transcriptional regulator (H0N27_10825; TR), we aimed to understand its role in the identified system. Given its putative function, we first checked the effect of deleting the TR gene on the transcript levels of the RE and the MT during exponential growth (2h growth post-dilution). Interestingly, deletion of the TR gene resulted in a ~50-fold increase of the MT gene expression (Fig. 5A). No significant differences in RE transcript levels were detected in the ΔTR strain compared to its parental WT strain (Fig. 5A). These results indicated that the TR likely serves as a repressor of the MT gene, suggesting that the absence of this regulator could result in enhanced levels of methylation. To test this, we assessed the DNA methylome of the TR-deficient strain by SMRT sequencing. Interestingly, the DNA methylome changed significantly in the TR-deficient strain as compared to the WT(reSEQ). Several new DNA modifications appeared, all of which included a ‘GATC’ core sequence (Fig. 5B). These results suggest that the TR controls both the abundance and as a result, the specificity of the methylase.

**Figure 5.**
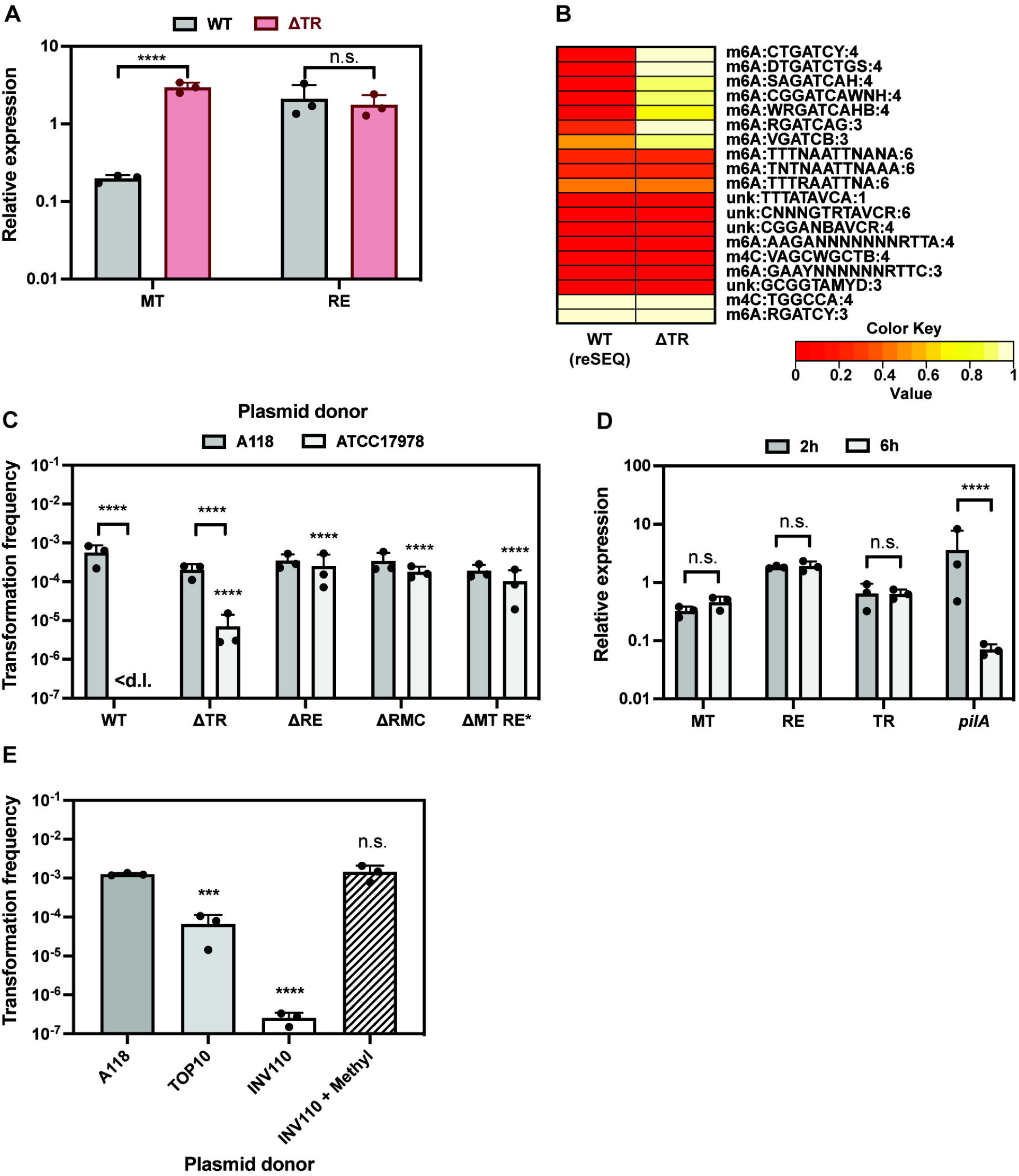
TR deficiency can overcome DNA restriction. **(A)** Relative expression of the MT and the RE genes in the TR-deficient mutant strain (ΔTR) compared to the parental WT strain A118. **(B)** Heatmap showing the abundance of different DNA modifications in WT and ΔTR. Details as described for Figure 2. **(C)** Transformability of strain A118 and its RMC-related derivatives using self and non-self plasmid DNA (isolated from A118 or ATCC17978) as transforming material. Transformation frequencies of RMC-related derivatives are compared to the WT within each set that tested the same plasmid source. Only statistically significant differences are shown. **(D)** Relative expression of the *MT, RE, TR*, and *pilA* genes in strain A118 after 2h and 6h of growth. **(E)**Transformation frequency of strain A118 with plasmids pABA originating from strains A118, *E. coli* TOP10 and *E. coli* INV110. Dam methylase-dependent *in vitro* methylation was performed where indicated (+ Methyl). The mean (± SD) was calculated from three independent biological replicates (A, C-E). <d.l., below detection limit. The data was log-transformed and statistical differences were calculated using two-way ANOVA corrected for multiple comparisons with Šidák’s method (A, C-D) or ordinary one-way ANOVA with Dunnett’s correction for multiple comparisons (E). ***, *P* < 0.001, ****, *P* < 0.0001; n.s., not significant.

Given these results, we hypothesized that over-abundance of the methylase might lead to inappropriate protection of incoming non-self DNA and therefore reduce the selectivity for self-DNA. We therefore compared the transformability of the TR-minus strain with the WT and several RE-deficient derivatives. As transforming material, we used either plasmid DNA or gDNA originating from strain A118 (‘self’) or ATCC17978 (‘non-self’). As expected, all strains were transformed equally well by self-DNA (Figs 5C and S4A). However, for the non-self plasmid DNA, we observed different transformation patterns depending on the acceptor strain. Indeed, the transformation frequency was under the detection limit in the WT strain but increased dramatically in all other strains, irrespective of those being RE-proficient or RE-deficient (Fig. 5C). Strikingly, transformation by the nonself plasmid was also clearly detectable for the TR-minus strain (Fig. 5C). A similar pattern was also observed when non-self gDNA from ATCC17978 was used as transforming material, with intermediate transformation levels for the TR-deficient strain (Fig. S4A). Taken together, these results suggest that overabundance of the methylase can result in (partial) protection of foreign DNA.

To confirm that the observed phenotype is caused by the TR, we performed transformation assays with the WT, ΔTR strain, and a complemented version, wherein the TR-encoding gene (H0N27_10825) was placed under the control of its native promoter at the *hcp* locus, along with a gentamicin-resistant cassette (GentR) (for selection during strain construction). The resulting strains were tested for their transformability using foreign plasmid DNA derived from strain ATCC17978. Consistent with the results above, transformation frequencies were close to the detection limit for the WT control strain that carried the GentR cassette within *hcp* (Fig. S4B). However, in the absence of the TR, transformants were readily detectable. This effect was reversed in the strain containing the complemented TR gene (Fig. S4B). These results confirm that overproduction of the methylase in the ΔTR strain was indeed caused by the absence of the TR and was not a result of a polar effect on the MT gene.

Given the observation that the MT transcription is tightly controlled, we examined whether the MT, RE, and TR levels vary during different growth phases. Indeed, since we previously observed that some of the essential competence genes – e.g., type IV pili (T4P) genes – are only produced during exponential growth (46), we wondered whether methylation levels are also increased specifically during this ‘transformation window’, potentially to facilitate the acquisition of new genetic traits from bacteria lacking ‘GATC’-like methylation marks. However, in contrast to the major T4P gene *pilA*, our results indicate that all the RMC genes are stably expressed in exponential and stationary growth phase (Fig. 5D). This suggests that A118 does not adapt its defense mechanism against foreign DNA during the transformation process. This in contrast to *S. pneumoniae*, for which the ssDNA methylase DpnA was shown to be specifically expressed during competence so that incoming transforming DNA was protected from degradation by an endogenous RM system (75, 81). Nevertheless, we cannot exclude that external factors might result in the increased MT levels in strain A118 under so far unknown conditions, which could facilitate the uptake of foreign DNA during high demand for new genetic traits, for instance, under harsh environmental conditions.

### *Escherichia coli* Dam methylation protects DNA from RMC-dependent degradation

Our data so far are consistent with the DNA modification m6A:RGATCY:3 being responsible for the DNA-origin-dependent transformability phenotype of strain A118 (Fig. 2A). Furthermore, the RE we have identified within the RMC belongs to the *Bgl*II/*Bst*YI endonuclease protein family, which generally degrades AGATCT or RGATCY. Interestingly, a common feature of these motifs is the ‘GATC’ core, a DNA sequence that is also recognized by the orphan methylase Dam (82). The Dam methylase was originally discovered in *E. coli* (82) and is present in many other Gamma-proteobacteria. It is, however, absent in the genus *Acinetobacter*. Dam recognizes GATC and adds a methyl group onto the adenine. We therefore hypothesized that Dam methylation could serve to protect non-self DNA during transformation of strain A118. To test this idea, we purified pABA from *E. coli* strains that were either *dam+* (TOP10) or *dam-* (INV110) and used them as donor DNA in transformation assays. Notably, pABA purified from the *dam*+ *E. coli* TOP10 supported efficient transformation (~10^-4^), whereas transformation was close to the detection limit (~10^-7^) when the plasmid was purified from the *dam-* strain INV110 (Fig. 5E). The transformation frequency remained around 10-fold lower for TOP10-derived plasmid DNA compared to self DNA from *A. baumannii* strain A118. One reason for this discrepancy could be the incomplete methylation of the plasmid, possibly due to the depletion of the methyl group donor (S-adenosyl-L-methionine, SAM) or an additional DNA modification acquired in TOP10 that negatively affects transformation. To test the former idea, we performed an *in vitro* methylation assay whereby we methylated plasmid DNA derived from *E. coli* strain INV110 with commercially available Dam methylase (New England BioLabs) and tested whether it affected plasmid transformation in strain A118. Strikingly, the *in vitro* methylation rendered the plasmid fully transformation-competent leading to transformation frequencies comparable to those obtained with A118-derived plasmid DNA (Fig. 5E). These results demonstrated that Dam methylation can efficiently protect DNA from degradation by the RMC-associated RE. Interestingly, according to REBASE restriction enzyme database (rebase.neb.com/rebase/) this is not an efficient protection against the commercially available restriction enzymes *Bgl*II and *Bst*YI, as Dam methylation does not have an inhibitory effect on their DNA digestion activity.

Taken together, these results strongly suggest that the RMC could play an important role in blocking DNA exchange from different bacteria lacking ‘GATC’-related methylation marks. However, our results also demonstrate that Dam methylation can bypass the RMC-dependent effect. Interestingly, according to the REBASE database, many ESKAPE pathogens, such as *Pseudomonas aeruginosa*, *Klebsiella pneumoniae* and *Enterobacter spp*., carry Dam or Dam-related methylation motifs, making them a possible reservoir of resistance genes for *A. baumannii* strains with the modules similar to the described RMC.

## CONCLUSION

In this study, we described a RM-dependent mechanism that significantly affects intra- and interspecific DNA exchange with competent *A. baumannii* strain A118. We showed that the DNA methylome is variable between the tested *A. baumannii* strains, which was partially reflected in the ability of the strains’ DNA to serve as efficient transforming material. Furthermore, we identified an A118-specific RM system, which fostered the degradation of foreign DNA that lacked specific epigenetic marks. This resulted in indirect enhancement of transformation with kin-released DNA, suggesting that the presence of the RM systems in competent cells can reduce the level of transformation-driven genome diversification. Indeed, for strain A118, transforming DNA derived from foreign strains will be less efficiently incorporated compared to self-DNA thereby likely reducing the acquisition of novel genetic traits. While DNA acquisition is considered one of the primary roles of natural transformation (64), another role of transformation, namely genome curing from mobile genetic elements (83), might remain highly efficient. The primary role of RM systems is considered to be the protection against invasion of plasmids and bacteriophages (84). Interestingly, the studied RM system was found in a genomic region that varied significantly between different *A. baumannii* strains, but ubiquitously carried genes involved in DNA defense including anti-phage systems. The effect of the identified RM system on transformation might therefore be a consequence of anti-phage or anti-plasmid strategies, originally aiming to prevent phage infection and preserve genome content.

However, our data suggested that the RM-dependent effect on transformation could be avoided to some extent when the level of the methylase was increased in the cell. Speculatively, this mechanism could facilitate the uptake of foreign DNA under stressful environmental conditions when new genetic features could serve as a bet-hedging strategy enabling bacterial survival. Additionally, transforming DNA carrying overlapping Dam methylation marks was not impacted by the presence of the RM system suggesting a preference for DNA originating from bacterial species that carry similar epigenetic marks.

These findings could foster future research that aims at understanding the reservoir of horizontally-acquired DNA fragments, including the frequently observed chromosomally-located AMR genes of diverse *A. baumannii* isolates. Since *A. baumannii* is one of the most threatening multidrug resistant pathogens, it is crucial to understand how it acquires dangerous genetic traits.

## Data availability

The assembled genome sequences and raw data were deposited into NCBI. GenBank accession numbers and BioSample IDs are indicated in the Tables S2 and S3.

## Supplementary data

Supplementary data are available at XXXX online.

## Acknowledgements

We thank Tu Giang Doan for technical assistance, X. Charpentier for sharing the published *A. baumannii* isolates, and members of Blokesch lab, in particular David. W. Adams and Nicolas Flaugnatti for fruitful discussions. We also thank the staff of the Lausanne Genomic Technologies Facility at the University of Lausanne for sample processing, genome sequencing, and basic bioinformatic analysis.

## Author contributions

M.B. secured funding and supervised the study; M.B and N.V conceived the study and designed the experiments; N.V. performed the wetlab experiments; C.I. and N.G. performed bioinformatic analyses to identify DNA modifications; A.L. performed bioinformatic analyses for genome comparison of *A. baumannii* strains; N.V. and M.B. analyzed the data and wrote the manuscript with input from C.I., N.G., and A.L. All authors approved the final version of the manuscript.

