## Supplementary material for "DNA modifications impact natural transformation of *Acinetobacter baumannii*"

**Table S1. Bacterial strains and plasmids used in this study**

| Strains or plasmids | Genotype/description* | Internal strain number | Reference (original strain and genome sequence) |
|---|---|---|---|
| ***A. baumannii*** | | | |
| A118 | Wild type; $Amp^R$, $Cm^R$ | MB#5144 | (1–3) |
| ATCC17978 | Wild type; $Amp^R$, $Cm^R$ | MB#5143 | (4) |
| ATCC19606 | Wild type; $Amp^R$, $Cm^R$, $Strep^R$ | MB#5145 | (5–7) |
| AB5075 | Wild type; $Amp^R$, $Kan^R$ | MB#8577 | (8–10) |
| 29D2 | Wild type; $Amp^R$ | MB#8581 | (11) |
| 86II/2C | Wild type; $Amp^R$ | MB#8582 | (11) |
| A118Δhcp::aprR | A118 with *hcp* replaced by *aac(3)IV* cassette using natural transformation; $Amp^R$, $Cm^R$, $Apr^R$ | MB#10119 | This study |
| ATCC17978Δhcp::aprR | ATCC17978 with *hcp* replaced by *aac(3)IV* cassette using electroporation; $Amp^R$, $Cm^R$, $Apr^R$ | MB#10120 | This study |
| ATCC19606Δhcp::aprR | ATCC19606 with *hcp* replaced by aac(3)IV cassette using suicide plasmid pGP704Sac28-Δhcp::aprR; $Amp^R$, $Cm^R$, $Strep^R$, $Apr^R$ | MB#10121 | This study |
| AB5075Δhcp::aprR | AB5075 with *hcp* replaced by *aac(3)IV* cassette using natural transformation; $Amp^R$, $Cm^R$, $Kan^R$, $Apr^R$ | MB#10122 | This study |
| 29D2Δhcp::aprR | 29D2 with *hcp* replaced by *aac(3)IV* cassette using natural transformation; $Amp^R$, $Apr^R$ | MB#10123 | This study |
| 86II/2CΔhcp::aprR | 86II/2C with *hcp* replaced by *aac(3)IV* cassette using natural transformation; $Amp^R$, $Apr^R$ | MB#10124 | This study |
| A118 / pABA | A118 carrying pABA; $Amp^R$, $Cm^R$, $Apr^R$ | MB#10125 | This study |
| ATCC17978 / pABA | ATCC17978 carrying pABA; $Amp^R$, $Cm^R$, $Apr^R$ | MB#10126 | This study |
| ATCC19606 / pABA | ATCC19606 carrying pABA; $Amp^R$, $Cm^R$, $Strep^R$, $Apr^R$ | MB#10127 | This study |
| AB5075 / pABA | AB5075 carrying pABA; $Amp^R$, $Cm^R$, $Kan^R$ ,$Apr^R$ | MB#10128 | This study |
| 29D2 / pABA | 29D2 carrying pABA; $Amp^R$, $Apr^R$ | MB#10129 | This study |
| 86II/2C / pABA | 86II/2C carrying pABA; $Amp^R$, $Apr^R$ | MB#10130 | This study |
| A118ΔH0N27_10830::kanR (ΔRE) | A118 with *H0N27_10830* deleted by insertion of *aph* cassette; $Amp^R$, $Cm^R$, $Kan^R$ | MB#9992 | This study |
| A118ΔH0N27_10820-30::kanR (ΔRMC) | A118 with *H0N27_10820-30* deleted by insertion of *aph* cassette; $Amp^R$, $Cm^R$, $Kan^R$ | MB#9993 | This study |
| A118ΔH0N27_10820-30::kanR / pABA (ΔRMC / pABA) | A118ΔH0N27_10820-30::kanR carrying pABA; $Amp^R$, $Cm^R$, $Kan^R$, $Apr^R$ | MB10131 | This study |
| A118ΔH0N27_10820-30::kanR-TnAraC (ΔRMC-Tn) | A118ΔH0N27_10820-30::kanR containing mini-Tn7-*araC* (TnAraC); $Amp^R$, $Cm^R$, $Kan^R$, $Gent^R$ | MB#10113 | This study |
| A118ΔH0N27_10820-30::kanR-TnAraC / pABA (ΔRMC-Tn / pABA) | A118ΔH0N27_10820-30::kanR-TnAraC carrying pABA; $Amp^R$, $Cm^R$, $Kan^R$, $Gent^R$, $Apr^R$ | MB#10132 | This study |

| | | | |
|---|---|---|---|
| A118ΔH0N27_10820-30::kanR-TnH0N27_10820 (ΔRMC-TnMT) | A118ΔH0N27_10820-30::kanR containing mini-Tn7-*araC-H0N27_10820* (TnH0N27_10820); $Amp^R$, $Cm^R$, $Kan^R$, $Gent^R$ | MB#10114 | This study |
| A118ΔH0N27_10820-30::kanR-TnH0N27_10820 / pABA (ΔRMC-TnMT / pABA) | A118ΔH0N27_10820-30::kanR TnH0N27_10820 carrying pABA; $Amp^R$, $Cm^R$, $Kan^R$, $Gent^R$, $Apr^R$ | MB#10133 | This study |
| A118ΔH0N27_10820::kanR-H0N27_10830-K176N (ΔMT RE*) | A118 with *H0N27_10820* deleted by insertion of *aph* cassette plus spontaneous mutation in H0N27_10830; $Amp^R$, $Cm^R$, $Kan^R$ | MB#9995 | This study |
| A118ΔH0N27_10820::kanR-H0N27_10830-K176N / pABA (ΔMT RE* / pABA) | A118ΔH0N27_10820::kanR H0N27_10830-K176N carrying pABA; $Amp^R$, $Cm^R$, $Kan^R$, $Gent^R$, $Apr^R$ | MB#10134 | This study |
| A118ΔH0N27_10820::kanR-H0N27_10830-K176N-TnAraC (ΔMT RE*-Tn) | A118ΔH0N27_10820::kanR H0N27_10830-K176N containing mini-Tn7-*araC* (TnAraC); $Amp^R$, $Cm^R$, $Kan^R$, $Gent^R$ | MB#10115 | This study |
| A118ΔH0N27_10820::kanR-H0N27_10830-K176N-TnAraC / pABA (ΔMT RE*-Tn / pABA) | A118ΔH0N27_10820::kanR H0N27_10830-K176N TnAraC carrying pABA; $Amp^R$, $Cm^R$, $Kan^R$, $Gent^R$, $Apr^R$ | MB#10135 | This study |
| A118ΔH0N27_10820::kanR-H0N27_10830-K176N-TnH0N27_10820 (ΔMT RE*-TnMT) | A118ΔH0N27_10820::kanR H0N27_10830-K176N containing mini-Tn7-*araC-H0N27_10820* (TnH0N27_10820); $Amp^R$, $Cm^R$, $Kan^R$, $Gent^R$ | MB#10116 | This study |
| A118ΔH0N27_10820::kanR-H0N27_10830-K176N-TnH0N27_10820 / pABA (ΔMT RE*-TnMT / pABA) | A118ΔH0N27_10820::kanR H0N27_10830-K176N-TnH0N27_10820 carrying pABA; $Amp^R$, $Cm^R$, $Kan^R$, $Gent^R$, $Apr^R$ | MB#10136 | This study |
| ATCC17978-TnAraC (ATCC17978-Tn) | ATCC17978 containing mini-Tn7-*araC* (TnAraC); $Amp^R$, $Cm^R$, $Gent^R$ | MB#10117 | This study |
| ATCC17978-TnAraC / pABA (ATCC17978-Tn / pABA) | ATCC17978-TnAraC carrying pABA; $Amp^R$, $Cm^R$, $Gent^R$, $Apr^R$ | MB#10137 | This study |
| ATCC17978-TnH0N27_10820 (ATCC17978-TnMT) | ATCC17978 containing mini-Tn7-*araC-H0N27_10820* (TnH0N27_10820); $Amp^R$, $Cm^R$, $Gent^R$ | MB#10118 | This study |
| ATCC17978-TnH0N27_10820 / pABA (ATCC17978 TnMT / pABA) | ATCC17978 TnH0N27_10820 carrying pABA; $Amp^R$, $Cm^R$, $Gent^R$, $Apr^R$ | MB#10138 | This study |
| ATCC17978-TnH0N27_12600 (ATCC17978-TnMT-2) | ATCC17978 containing mini-Tn7-*araC-H0N27_12600* (TnH0N27_12600); $Amp^R$, $Cm^R$, $Gent^R$ | MB#10003 | This study |
| ATCC17978-TnH0N27_12600 / pABA (ATCC17978-TnMT-2 / pABA) | ATCC17978-TnH0N27_12600 carrying pABA; $Amp^R$, $Cm^R$, $Gent^R$, $Apr^R$ | MB#10139 | This study |
| A118ΔH0N27_10825::kanR (ΔTR) | A118 with *H0N27_10825* deleted by insertion of *aph* cassette using natural transformation; $Amp^R$, $Cm^R$, $Kan^R$ | MB#9994 | This study |
| A118ΔH0N27_10825::kanR / pABA (ΔTR / pABA) | A118ΔH0N27_10825::kanR carrying pABA, $Amp^R$, $Cm^R$, $Kan^R$, $Apr^R$ | MB#10140 | This study |
| A118Δhcp::gentR | A118 with *hcp* replaced by *aacC1* cassette using natural transformation; $Amp^R$, $Cm^R$, $Gent^R$ | MB#10141 | This study |
| A118ΔH0N27_10825::kanR Δhcp::gentR | A118ΔH0N27_10825::kanR with *hcp* replaced by *aacC1* cassette using natural transformation; $Amp^R$, $Cm^R$, $Gent^R$ | MB#10142 | This study |
| A118ΔH0N27_10825::kanR Δhcp::H0N27_10825-gentR | A118ΔH0N27_10825::kanR with *hcp* replaced by *H0N27_10825* & *aacC1* cassette using natural transformation; $Amp^R$, $Cm^R$, $Gent^R$ | MB#10143 | This study |

| | | | |
|---|---|---|---|
| ***E. coli*** | | | |
| S17-1λpir | $Tp^R$ $Sm^R$ *recA, thi, pro, hsdR2M1*, RP4:2-Tc:Mu:$Km^R$, Tn7 (λpir) | MB#648 | (12) |
| TOP10 | F- *mcrA* Δ( *mrr-hsd*RMS-*mcr*BC) Φ80*lac*ZΔM15 Δ *lac*X74 *rec*A1 *ara*D139 Δ( *araleu*)7697 *gal*U *gal*K *rps*L ($Strep^R$) *end*A1 *nup*G | MB#4817 | Invitrogen |
| INV110 | F´ {traΔ36 proAB lacIq lacZΔM15} rpsL ($Strep^R$) thr leu endA thi-1 lacY galK galT ara tonA tsx dam dcm supE44 Δ(lac-proAB) Δ(mcrC-mrr)102::Tn10 ($Tet^R$) | MB#9225 | Invitrogen |
| **Plasmids** | | | |
| pGP704-TnAraC | pGP704 with mini-Tn7 carrying *araC* and $P_{BAD}$; $Amp^R$, $Gent^R$ | MB#5513 | (13) |
| pGP704-TnH0N27_10820 | pGP704 with mini-Tn7 carrying *araC* and $P_{BAD}$- driven *H0N27_10820*; $Amp^R$, $Gent^R$ | MB#10144 | This study |
| pGP704-TnH0N27_12600 | pGP704 with mini-Tn7 carrying *araC* and $P_{BAD}$- driven *H0N27_12600*; $Amp^R$, $Gent^R$ | MB#10145 | This study |
| pUX-BF-13 | *ori*R6K, helper plasmid with Tn7 transposition function; $Amp^R$ | MB#457 | (14) |
| pGP704-Sac28 | suicide plasmid, ori R6K sacB; $Amp^R$ | MB#649 | (15) |
| pGP704-Sac28-Δhcp::aprR | pGP704Sac28 carrying a deletion within *hcp* and insertion of *aac(3)IV* cassette; $Amp^R$, $Apr^R$ | MB#10146 | This study |
| pABA | Derivative of pBAD18-Kan, with added *Acinetobacter*-specific ori (16), and replacement of *aph* cassette (KanR) for *aac(3)IV* cassette (AprR) | MB#9356 | This study |

*Locus tags of strain A118 and annotations are according to NCBI accession number CP059039.

### Table S2. Statistics of PacBio SMRT genome sequencing and assembly for WT strains

| | ATCC17978 | ATCC19606 | AB5075 | 29D2 | 86II/2C |
|---|---|---|---|---|---|
| **Internal strain ID** | MB#5143 | MB#5145 | MB#8577 | MB#8581 | MB#8582 |
| **BioSample ID** | SAMN15507634 | SAMN15507635 | SAMN31681936 | SAMN31681934 | SAMN31681935 |
| **GenBank accession number** | CP059041 | CP059040 | CP113078-CP113079 | CP113069 | CP113077 |
| **Chromosome length after circularization** | 4,006,609 | 3,980,852 | 3,995,103 | 3,836,532 | 3,937,483 |
| **Total genome size** | 4,006,609 | 3,980,852 | 4,078,713 | 3,836,532 | 3,937,483 |
| **Mean coverage** | 399.24 | 217.27 | 529.45 | 257 | 403.04 |
| **GC content** | 40.90 | 40.20 | 38.91 | 38.91 | 38.90 |

**Table S3. Statistics of PacBio SMRT genome sequencing and assembly mutant strains of A118 or ATCC17978**

| | ΔME RE* | ΔTR | ΔRE | ΔRMC | ATCC17978-TnMT-2 |
|---|---|---|---|---|---|
| **Internal strain ID** | MB#9995 | MB#9994 | MB#9992 | MB#9993 | MB #10003 |
| **BioSample ID** | SAMN31681941 | SAMN31681938 | SAMN31681939 | SAMN31681940 | SAMN31681942 |
| **GenBank accession number** | CP113073 | CP113070 | CP113071 | CP113072 | CP113074-CP113076 |
| **Chromosome length after circularization** | 3,750,381 | 3,751,367 | 3,750,940 | 3,749,418 | 4,017,814 |
| **Total genome size** | 3,750,381 | 3,751,367 | 3,750,940 | 3,749,418 | 4,042,525 |
| **Mean coverage** | 468.94 | 455.43 | 465.99 | 342.69 | 420.03 |
| **GC content** | 39.42 | 39.43 | 39.49 | 39.40 | 39.34 |

**Table S4. Characteristics of *A. baumannii* strains**

| Strains name | Year of isolation | Country of isolation | Host | Sample type | Naturally transformable | Reference |
|---|---|---|---|---|---|---|
| A118 | 1995 | Argentina | Human | Blood | yes | (1, 2) |
| ATCC17978 | 1951 | ? | Human | Spinal meningitis | no | (4, 17) |
| ATCC19606 | 1948 | US | Human | Urine | no | (5–7) |
| 29D2 | 2014 | Poland | White stork | Nestling; choana | yes | (11) |
| 86II/2C | 2013 | Poland | White stork | Nestling; choana | yes | (11) |
| AB5075 | 2008 | US | Human | Tibia/ osteomyelitis | yes | (8, 10) |

**Table S5. Number of identified recognition in analyzed *A. baumannii* genomes, plasmids, and PCR fragments**

| Epigenetic mark | A118 | ATCC 17978 | ATCC 19606 | AB507 5 | 29D2 | 86II/2C | plasmid pABA | PCR Δ*hcp*:: AprR | PCR *rpoB** |
|---|---|---|---|---|---|---|---|---|---|
| unk:CNNNGTRTAVCR:6 | 143 | 149 | 160 | 152 | 148 | 155 | 0 | 0 | 0 |
| unk:CGGANBAVCR:4 | 192 | 222 | 219 | 205 | 203 | 205 | 2 | 1 | 0 |
| unk:CGGTGTNTY:4 | 144 | 159 | 172 | 154 | 152 | 170 | 0 | 0 | 0 |
| unk:TTTATAVCA:1 | 177 | 173 | 176 | 191 | 175 | 173 | 1 | 0 | 1 |
| m6A:TTTRAATTNA:6 | 519 | 578 | 550 | 545 | 532 | 526 | 0 | 1 | 0 |
| m6A:TNTNAATTNAAA:6 | 374 | 386 | 365 | 366 | 356 | 364 | 0 | 2 | 0 |
| m6A:TAAYNNNNNNNTCTT:3 | 567 | 600 | 581 | 588 | 577 | 585 | 2 | 0 | 1 |
| m6A:CTATCAV:6 | 1096 | 1214 | 1202 | 1217 | 1129 | 1156 | 3 | 2 | 2 |
| m6A:AAGANNNNNNNRTTA:4 | 567 | 600 | 581 | 588 | 577 | 585 | 2 | 0 | 1 |
| unk:GCGGTAMYD:3 | 313 | 327 | 322 | 327 | 306 | 330 | 1 | 0 | 0 |
| m6A:RGATCY:3 | 2842 | 3404 | 3150 | 3294 | 3090 | 3080 | 11 | 4 | 1 |
| unk:TGGCCA:4 | 1264 | 1398 | 1374 | 1418 | 1352 | 1316 | 0 | 0 | 0 |
| m6A:GAAYNNNNNNRTTC:3 | 518 | 574 | 540 | 554 | 518 | 512 | 0 | 1 | 1 |
| m4C:VAGCWGCTB:4 | 1414 | 1456 | 1508 | 1528 | 1422 | 1408 | 2 | 0 | 2 |
| m6A:GAAAGC:4 | 2131 | 2284 | 2306 | 2320 | 2175 | 2257 | 6 | 1 | 2 |
| m6A:CTGATCY:4 | 744 | 822 | 832 | 805 | 751 | 787 | 3 | 1 | 0 |
| m6A:DTGATCTGS:4 | 193 | 219 | 215 | 227 | 218 | 216 | 1 | 1 | 0 |
| m6A:SAGATCAH:4 | 557 | 647 | 619 | 633 | 595 | 619 | 2 | 1 | 0 |
| m6A:CGGATCAWNH:4 | 177 | 170 | 172 | 158 | 187 | 183 | 2 | 1 | 0 |
| m6A:WRGATCAHB:4 | 1376 | 1488 | 1471 | 1496 | 1385 | 1462 | 1 | 2 | 0 |
| m6A:RGATCAG:3 | 744 | 822 | 832 | 805 | 751 | 787 | 3 | 1 | 0 |
| m6A:VGATCB:3 | 6288 | 7210 | 6840 | 7178 | 6658 | 6688 | 15 | 8 | 3 |
| m6A:TTTNAATTNANA:6 | 374 | 386 | 365 | 366 | 356 | 364 | 0 | 2 | 0 |

### Table S6. BLAST results of the RMC genomic region

| Description | Scientific Name | Max Score | Total Score | Query Cover | E value | Per. ident | Acc. Len | Accession |
|---|---|---|---|---|---|---|---|---|
| ***Acinetobacter baumannii* strain A118 chromosome, complete genome (input query)** | *Acinetobacter baumannii* | 3991 | 3991 | 100% | 0.0 | 100.00 | 3750370 | NZ_CP059039.1 |
| ***Acinetobacter baumannii* strain A118F NODE_11_length_136216_cov_13.6381, whole genome shotgun sequence** | *Acinetobacter baumannii* | 3991 | 3991 | 100% | 0.0 | 100.00 | 136216 | NZ_VCCO01000013.1 |
| ***Acinetobacter baumannii* strain Ab42 NODE_1_length_802719_cov_1.259005, whole genome shotgun sequence** | *Acinetobacter baumannii* | 3991 | 3991 | 100% | 0.0 | 100.00 | 802719 | NZ_JAAZUE010000001.1 |
| ***Acinetobacter baumannii* strain ARLG-1772 PR391_NODE_5.ctg_1, whole genome shotgun sequence** | *Acinetobacter baumannii* | 3980 | 3980 | 100% | 0.0 | 99.91 | 245745 | NZ_NGEY01000012.1 |
| ***Acinetobacter baumannii* UH6507 ctgN19, whole genome shotgun sequence** | *Acinetobacter baumannii* UH6507 | 3980 | 3980 | 100% | 0.0 | 99.91 | 313343 | NZ_AYFK01000017.1 |
| ***Acinetobacter oleivorans* strain KCJK7897 NODE_3_length_378121_cov_27.2151_ID_5, whole genome shotgun sequence** | *Acinetobacter oleivorans* | 3105 | 3105 | 100% | 0.0 | 92.60 | 378121 | NZ_QAYN01000006.1 |
| ***Acinetobacter baumannii* strain MRSN32076 MRSN32076_MRSN32076_contig00061, whole genome shotgun sequence** | *Acinetobacter baumannii* | 2850 | 2850 | 100% | 0.0 | 90.47 | 32390 | NZ_VHFM01000061.1 |
| ***Acinetobacter baumannii* NIPH 60 acLrl-supercont1.29, whole genome shotgun sequence** | *Acinetobacter baumannii* NIPH 60 | 2850 | 2850 | 100% | 0.0 | 90.47 | 494539 | NZ_KB849502.1 |
| ***Acinetobacter baumannii* strain ABBL038 contig-37, whole genome shotgun sequence** | *Acinetobacter baumannii* | 2844 | 2844 | 100% | 0.0 | 90.43 | 95751 | NZ_LLDS01000058.1 |
| ***Acinetobacter baumannii* strain ABBL110 contig-3000004, whole genome shotgun sequence** | *Acinetobacter baumannii* | 1701 | 1701 | 57% | 0.0 | 91.27 | 8035 | NZ_LLHI01000148.1 |
| ***Acinetobacter tandoii* strain SC36 contig39, whole genome shotgun sequence** | *Acinetobacter tandoii* | 1107 | 1107 | 97% | 0.0 | 76.48 | 11215 | NZ_LBNL01000039.1 |
| ***Acinetobacter* sp. YH12083 NODE_27_length_6361_cov_74.148736, whole genome shotgun sequence** | *Acinetobacter* sp. YH12083 | 1075 | 1075 | 100% | 0.0 | 76.03 | 6361 | NZ_VPAD01000030.1 |
| ***Acinetobacter* sp. YH12133 NODE_29_length_6480_cov_156.757595, whole genome shotgun sequence** | *Acinetobacter* sp. YH12133 | 1075 | 1075 | 100% | 0.0 | 76.03 | 6480 | NZ_VPBS01000029.1 |
| ***Acinetobacter* sp. YH12139 NODE_29_length_6480_cov_184.474107, whole genome shotgun sequence** | *Acinetobacter* sp. YH12139 | 1075 | 1075 | 100% | 0.0 | 76.03 | 6480 | NZ_VPBY01000029.1 |
| ***Acinetobacter* sp. YH12132 NODE_29_length_6480_cov_145.652133, whole genome shotgun sequence** | *Acinetobacter* sp. YH12132 | 1075 | 1075 | 100% | 0.0 | 76.03 | 6480 | NZ_VPBR01000029.1 |
| ***Acinetobacter indicus* strain AI31 NODE_58_length_8886_cov_1445.717675, whole genome shotgun sequence** | *Acinetobacter indicus* | 1072 | 1072 | 100% | 0.0 | 75.98 | 8886 | NZ_JAAZRK010000058.1 |
| ***Acinetobacter indicus* strain AI29 NODE_57_length_8886_cov_817.336361, whole genome shotgun sequence** | *Acinetobacter indicus* | 1072 | 1072 | 100% | 0.0 | 75.99 | 8886 | NZ_JAAZRM010000057.1 |
| ***Acinetobacter* sp. YH12046 NODE_51_length_8267_cov_43.768796, whole genome shotgun sequence** | *Acinetobacter* sp. YH12046 | 1072 | 1072 | 100% | 0.0 | 75.98 | 8267 | NZ_VOZF01000052.1 |
| ***Acinetobacter* sp. YH12091 NODE_34_length_8267_cov_66.187715, whole genome shotgun sequence** | *Acinetobacter* sp. YH12091 | 1072 | 1072 | 100% | 0.0 | 75.99 | 8267 | NZ_VPAJ01000034.1 |

| | | | | | | | | |
|---|---|---|---|---|---|---|---|---|
| ***Acinetobacter*** **sp. YH12112 NODE_62_length_8267_cov_98.356880, whole genome shotgun sequence** | *Acinetobacter* sp. YH12112 | 1072 | 1072 | 100% | 0.0 | 75.99 | 8267 | NZ_VPBA01000063.1 |
| ***Acinetobacter*** **sp. YH12090 NODE_35_length_8267_cov_103.266830, whole genome shotgun sequence** | *Acinetobacter* sp. YH12090 | 1072 | 1072 | 100% | 0.0 | 75.98 | 8267 | NZ_VPAI01000035.1 |
| ***Acinetobacter*** **sp. YH1901152 NODE_54_length_8267_cov_61.999017, whole genome shotgun sequence** | *Acinetobacter* sp. YH1901152 | 1072 | 1072 | 100% | 0.0 | 75.99 | 8267 | NZ_VPFA01000056.1 |
| ***Acinetobacter indicus*** **strain AI21 NODE_57_length_6209_cov_1859.771200, whole genome shotgun sequence** | *Acinetobacter indicus* | 1018 | 1018 | 98% | 0.0 | 75.73 | 6209 | NZ_JAAZRU010000057.1 |
| ***Acinetobacter*** **sp. ANC 5347 Contig37, whole genome shotgun sequence** | *Acinetobacter* sp. ANC 5347 | 1000 | 1000 | 100% | 0.0 | 75.48 | 6481 | NZ_PGOZ01000037.1 |
| ***Acinetobacter lwoffii*** **strain VE243-2 contig00045, whole genome shotgun sequence** | *Acinetobacter lwoffii* | 929 | 929 | 84% | 0.0 | 76.17 | 6909 | NZ_JAMXXR010000045.1 |
| ***Acinetobacter*** **sp. YH12066 NODE_59_length_13185_cov_94.117016, whole genome shotgun sequence** | *Acinetobacter* sp. YH12066 | 854 | 854 | 83% | 0.0 | 75.53 | 13185 | NZ_VOZQ01000059.1 |
| ***Acinetobacter*** **sp. TG19627 A_sp_TG19627_7, whole genome shotgun sequence** | *Acinetobacter* sp. TG19627 | 843 | 843 | 83% | 0.0 | 75.42 | 7121 | NZ_AMJM01000006.1 |
| ***Acinetobacter genomosp.*** **16BJ strain CIP 70.18 acLso-supercont1.5, whole genome shotgun sequence** | *Acinetobacter genomosp.* 16BJ | 843 | 843 | 83% | 0.0 | 75.42 | 7141 | NZ_KB850080.1 |
| ***Acinetobacter*** **sp. YH16032 NODE_53_length_7154_cov_1106.629145, whole genome shotgun sequence** | *Acinetobacter* sp. YH16032 | 817 | 817 | 69% | 0.0 | 76.76 | 7154 | NZ_VPEE01000056.1 |
| ***Acinetobacter*** **sp. YH12105 NODE_37_length_6292_cov_57.114842, whole genome shotgun sequence** | *Acinetobacter* sp. YH12105 | 809 | 809 | 83% | 0.0 | 75.24 | 6292 | NZ_VPAU01000038.1 |
| ***Acinetobacter baumannii*** **strain ABBL110 contig-69, whole genome shotgun sequence** | *Acinetobacter baumannii* | 643 | 643 | 22% | 1E-180 | 90.39 | 14279 | NZ_LLHI01000265.1 |
| ***Acinetobacter*** **sp. YH01025 NODE_24_length_6480_cov_287.777743, whole genome shotgun sequence** | *Acinetobacter* sp. YH01025 | 505 | 505 | 37% | 6E-139 | 78.04 | 6480 | NZ_VOYR01000024.1 |

**A**

| | A118 | ATCC17978 | ATCC19606 | AB5075 | 29D2 | 86II/2C |
|---|---|---|---|---|---|---|
| A118 | 100 / 100 | 45.9 / 98.7 | 80.8 / 97.7 | 96.9 / 98.7 | 97.0 / 99.0 | 97.9 / 98.7 |
| ATCC17978 | | 100 / 100 | 41.7 / 97.6 | 45.8 / 98.8 | 46.1 / 98.9 | 46.1/ 99.0 |
| ATCC19606 | | | 100 / 100 | 79.9 / 97.6 | 80.4 / 97.7 | 80.7 / 97.8 |
| AB5075 | | | | 100 / 100 | 96.3 / 98.8 | 96.5 / 99.0 |
| 29D2 | | | | | 100 / 100 | 97.4 / 99.0 |
| 86II/2C | | | | | | 100 / 100 |

**B**

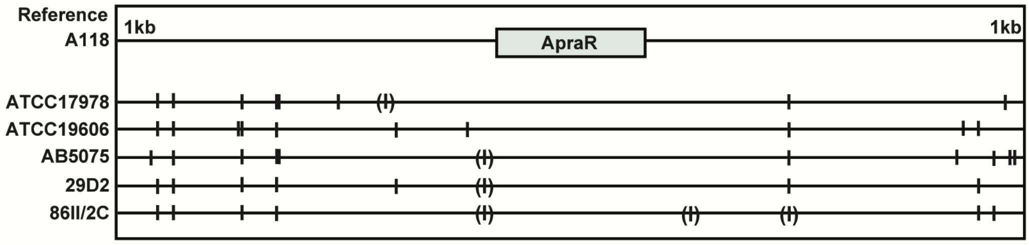


**Figure S1. Sequence identity surrounding the inserted apramycin cassette in different *A. baumannii* strains.**

**(A)** Pairwise identity of DNA sequences between different *A. baumannii* strains comparing 10-kb upstream / downstream from the inserted apramycin cassette. Values were calculated using the Geneious Prime software and are shown in percentage. **(B)** Alignment of DNA sequences from different *A. baumannii* strains depicting the 1-kb upstream and downstream region relative to the inserted apramycin cassette. Single nucleotide polymorphisms (SNPs) compared to the reference strain A118 are indicated by short vertical lines; if in brackets, the initial SNP was lost during the transformation-based strain construction.

| A118 | ATCC17978 | ATCC19606 | AB5075 | 29D2 | 86II2C | |
|---|---|---|---|---|---|---|
| H0N27_01045 | H0N28_17225 | H0N29_01055 | OSV63_08965 | OSV61_01025 | OSV60_01035 | class_I_SAM−dependent_methyltransferase |
| H0N27_01430 | H0N28_16775 | H0N29_01440 | OSV63_08575 | OSV61_01415 | OSV60_01560 | methyltransferase_domain−containing_protein |
| H0N27_01720 | H0N28_16485 | H0N29_01735 | OSV63_08285 | OSV61_01715 | OSV60_01850 | class_I_SAM−dependent_methyltransferase |
| H0N27_02900 | H0N28_15060 | H0N29_02925 | OSV63_06870 | OSV61_02900 | OSV60_03060 | SAM − dependent_methyltransferase |
| H0N27_03000 | H0N28_14960 | H0N29_03025 | OSV63_06770 | OSV61_03000 | OSV60_03160 | methyltransferase |
| H0N27_05920 | H0N28_11835 | H0N29_06195 | OSV63_02765 | OSV61_05980 | OSV60_06300 | class_I_SAM−dependent_methyltransferase |
| H0N27_09035 | H0N28_08710 | H0N29_09175 | OSV63_18685 | OSV61_09455 | OSV60_09715 | methyltransferase_domain−containing_protein |
| H0N27_10530 | H0N28_06940 | H0N29_10735 | OSV63_17215 | OSV61_10955 | OSV60_11240 | class_I_SAM−dependent_methyltransferase |
| H0N27_11860 | H0N28_05480 | H0N29_12680 | OSV63_15670 | OSV61_12340 | OSV60_12655 | class_I_SAM−dependent_methyltransferase |
| H0N27_14365 | H0N28_03290 | H0N29_15535 | OSV63_13355 | OSV61_14935 | OSV60_15010 | class_I_SAM−dependent_methyltransferase |
| H0N27_15025 | H0N28_02520 | H0N29_16195 | OSV63_12585 | OSV61_15585 | OSV60_16060 | class_I_SAM−dependent_methyltransferase |
| H0N27_15560 | H0N28_01985 | H0N29_16790 | OSV63_12045 | OSV61_16125 | OSV60_16650 | SAM−dependent_methyltransferase |
| H0N27_16335* | H0N28_01205* | H0N29_17560* | OSV63_11265* | OSV61_16905* | OSV60_17425* | DNA_adenine_methylase |
| H0N27_10820 | | | | | | DNA_adenine_methylase |
| H0N27_12600 | | | | | | DNA_adenine_methylase |
| | H0N28_08035 | | | | | methyltransferase_domain−containing_protein |
| | H0N28_10790 | | | | | DNA_cytosine_methyltransferase |
| | H0N28_14480 | H0N29_03505 | | | OSV60_03915 | methyltransferase_domain−containing_protein |
| | H0N28_05630 | H0N29_12525 | | | | DNA_cytosine_methyltransferase |
| | | H0N29_08515 | | | | DNA_cytosine_methyltransferase |
| | | H0N29_10075 | | | | class_I_SAM−dependent_DNA_methyltransferase |
| | | H0N29_11830 | | | | DNA_adenine_methylase |
| | | H0N29_12980 | | | | SAM−dependent_DNA_methyltransferase |
| | | | OSV63_06320 | | | N−6_DNA_methylase |
| | | | OSV63_09575 | | | DNA_cytosine_methyltransferase |
| | | | OSV63_14515 | OSV61_13800 | | site−specific_DNA−methyltransferase |
| | | | | | OSV60_07345 | DNA_cytosine_methyltransferase |
| | | | | | OSV60_07355 | site−specific_DNA−methyltransferase |
| | | | | | OSV60_13815 | class_I_SAM−dependent_DNA_methyltransferase |

**Figure S2. Annotated DNA methylases / methyltransferases in different *A. baumannii* strains.**

A table indicates the presence (gray) or absence (white) of a selection of DNA methylases or methyltransferases found in at least one of the six studied *A. baumannii* strains (shown below the table). The locus tags of the identified methylases / methyltransferases are indicated.

*, gene homologs to previously identified methylase-encoding *aamA* (49).

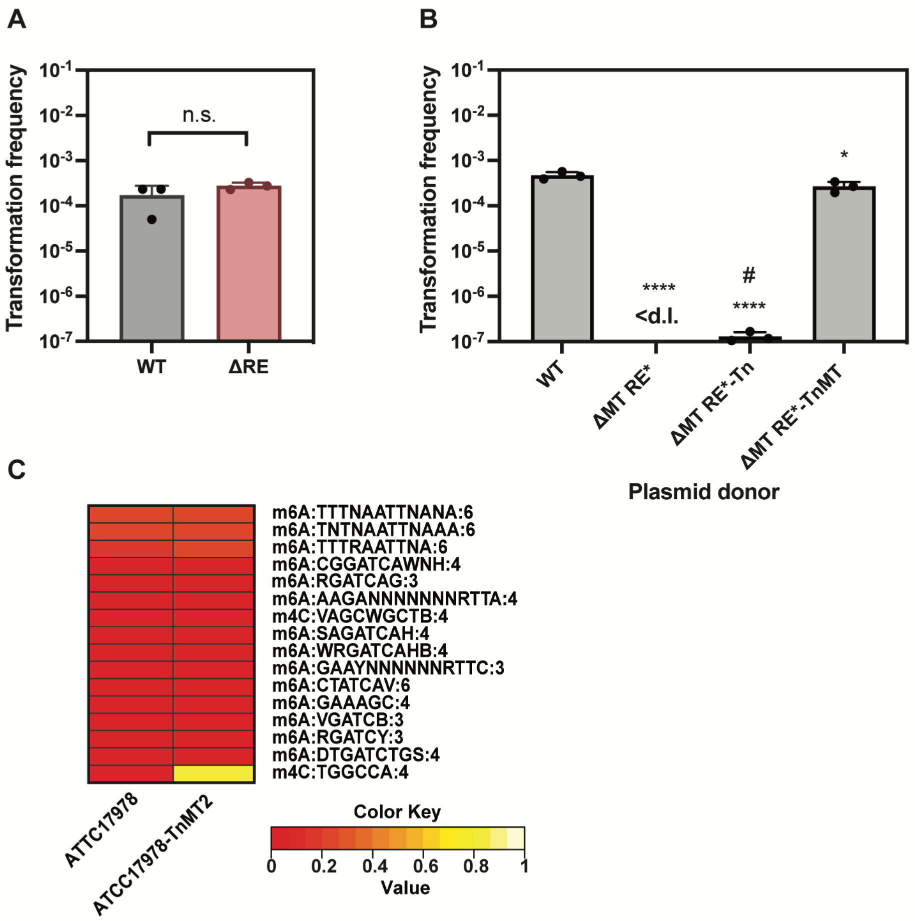


**Figure S3. RMC-related MT protects transforming DNA.**

**(A)** Transformability of strain A118 (WT) and A118ΔH0N27_10830 (ΔRE) using a PCR amplification product that encodes a rifampicin-resistant version of RpoB. **(B)** Transformability of strain A118 with plasmid pABA originating from various strains grown with 2% L-arabinose: A118 (WT), ΔH0N27_10820,H0N27_10830-K176N (ΔMT RE*), transposon control of ΔMT RE* (ΔMT RE*-Tn) or the H0N27_10820 (MT)-complemented version of ΔMT RE* (ΔMT RE*-TnMT). **(C)** Heatmap showing the abundance of certain DNA modification motifs in strain ATCC17978 and its MT2-overexpressing derivative (grown in presence of arabinose). Details as in Figure 2. The average (± SD) of three independent biological replicates is shown (A-B). <d.l., below detection limit. #, below detection limit in at least one replicate. The data was log-transformed and statistical differences were calculated using unpaired t-test (A) or a one-way ANOVA corrected for multiple comparisons with Dunnett's method (B). *, $P < 0.05$; **, $P < 0.01$; ****, $P < 0.0001$; n.s., not significant.

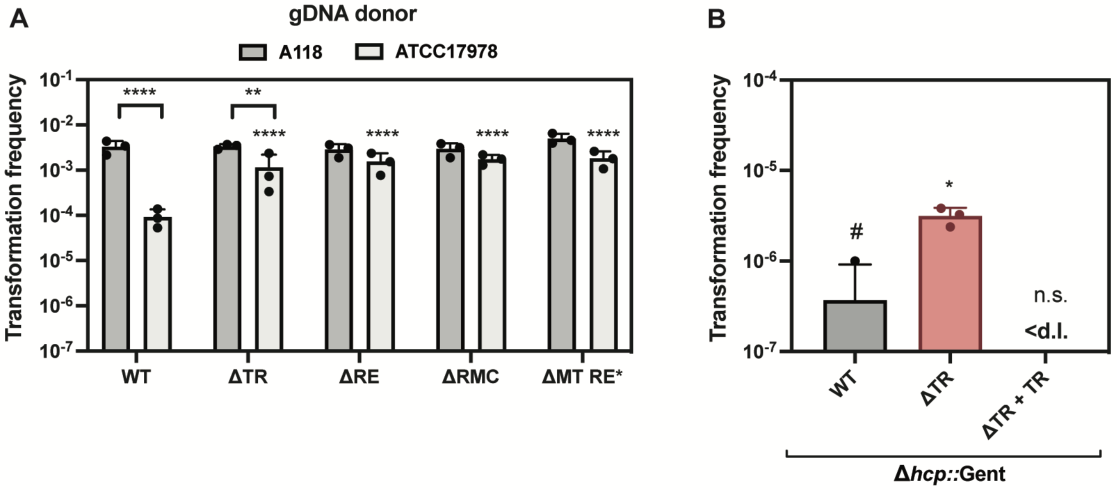


**Figure S4. Transformation with foreign DNA is facilitated in the TR-deficient strain.**

**(A)** Transformability of strain A118 and its RMC-related mutants using self (A118) and non-self (ATCC17978) genomic DNA as transforming material. Only statistically significant differences are shown. Samples for each strain after provision of different donor gDNA were compared. In addition, all strains within the specified gDNA sets were compared to the WT. **(B)** Transformability of A118 and its TR-related derivates using plasmid pABA isolated from strain ATCC17978 as transforming material. Complemented version of the TR-deficient mutant was constructed by addition of the TR gene inside *hcp* together with a gentamicin resistance cassette (ΔTR + TR). The WT and ΔTR contained the gentamicin resistance cassette inside *hcp* as a control. The mean (± SD) of three independent replicates is shown. <d.l., below detection limit. #, below detection limit in two replicates. The transformation frequency values were log-transformed and statistically analyzed using a two-way ANOVA corrected for multiple comparisons with Šidák's method (A) or an ordinary one-way ANOVA with Dunnett's correction for multiple comparisons (B). *, $P < 0.05$; **, $P < 0.01$; ****, $P < 0.0001$; n.s., not significant.
